# Distinct sets of molecular characteristics define tumor-rejecting neoantigens

**DOI:** 10.1101/2024.02.13.579546

**Authors:** Anngela C. Adams, Anne M. Macy, Elizabeth S. Borden, Lauren M. Herrmann, Chad A. Brambley, Tao Ma, Xing Li, Alysia Hughes, Denise J. Roe, Aaron R. Mangold, Kenneth H. Buetow, Melissa A. Wilson, Brian M. Baker, Karen Taraszka Hastings

## Abstract

Challenges in identifying tumor-rejecting neoantigens limit the efficacy of neoantigen vaccines to treat cancers, including cutaneous squamous cell carcinoma (cSCC). A minority of human cSCC tumors shared neoantigens, supporting the need for personalized vaccines. Using a UV-induced mouse cSCC model which recapitulated the mutational signature and driver mutations found in human disease, we found that CD8 T cells constrain cSCC. Two MHC class I neoantigens were identified that constrained cSCC growth. Compared to the wild-type peptides, one tumor-rejecting neoantigen exhibited improved MHC binding and the other had increased solvent accessibility of the mutated residue. Across known neoantigens that do not impact MHC binding, structural modeling of the peptide/MHC complexes indicated that increased solvent accessibility, which will facilitate TCR recognition of the neoantigen, distinguished tumor-rejecting from non-immunogenic neoantigens. This work reveals characteristics of tumor-rejecting neoantigens that may be of considerable importance in identifying optimal vaccine candidates in cSCC and other cancers.

## Main

Neoantigens are tumor-specific mutated peptides with the potential to elicit T cell-mediated tumor destruction.^1^ Neoantigen vaccines can improve clinical outcomes in patients with solid malignancies.^2, 3, 4, 5, 6^ However, the efficacy of neoantigen vaccines is limited by the challenge of selecting which neoantigen(s) mediate tumor rejection.^7^ Selection of which neoantigens to target is especially difficult in tumors with a high mutational burden, such as skin cancers which contain thousands of missense mutations. In high mutational burden tumors, neoantigens are often prioritized based on the prediction of improved MHC presentation, e.g., binding affinity and stability of the neoantigen/MHC complex and neoantigen expression. While this approach reliably identifies human and mouse immunogenic neoantigens, only a small percentage of prioritized neoantigens generate T cell responses.^8, 9, 10, 11, 12, 13, 14, 15, 16^ Furthermore, only a portion of the neoantigens which elicit T cell responses mediate tumor rejection.^9, 10, 11, 12, 17^ Thus, there is a need to evaluate the distinguishing characteristics of neoantigens which mediate tumor rejection.

Cutaneous squamous cell carcinoma (cSCC) is a high mutational burden tumor.^18^ Thus, cSCC is an ideal cancer to identify principles to improve the selection of neoantigens to target in neoantigen-based immunotherapies. To better understand characteristics of tumor-rejecting neoantigens, we developed a transplantable cSCC murine model from solar UV-induced tumors.^19^ This model recapitulates the mutational profile observed in human disease and has naturally occurring UV-induced neoantigens. The model is highly immunogenic and primarily constrained by CD8 T cells. Among the neoantigens with high predicted MHC class I binding affinity, MHC class I presentation, and expression, two tumor-rejecting neoantigens were identified. Compared to the wild-type peptides, one tumor-rejecting neoantigen exhibited improved MHC binding and the other had increased solvent accessibility of the mutated residue. For neoantigens that do not alter MHC binding, the predicted change in solvent accessibility of the mutated residue distinguished tumor-rejecting neoantigens in this and other studies from non-immunogenic neoantigens. This study highlights the addition of features anticipated to facilitate T cell receptor (TCR) recognition in the prioritization of vaccine candidates in cSCC as well as other cancers.

## Results

### Majority of mutations in human cSCC are unique to individual tumors

To assess the potential for shared, targetable neoantigens in cSCC, we analyzed the overlap of missense mutations in 149 human cSCC tumors from two datasets.^18, 20^ Compared to 139,276 mutations unique to a single tumor, there were only 27 mutations shared across four or more tumors and present in both datasets (Fig. 1a). As expected, since driver mutations are under positive selection in cancer,^21^ eight of the 27 shared mutations have been previously identified as putative driver mutations (Fig. 1a). The presence of shared mutations was not associated with sex or immune status (data not shown). Compared to tumors without shared mutations, tumors with shared mutations were from older patients (Extended Data Fig. 1a), patients less likely to have recessive dystrophic epidermolysis bullosa (RDEB), an inherited disease with increased risk of cSCC (Extended Data Fig. 1b), and had a higher mutational burden (Extended Data Fig. 1c). However, tumors from patients with RDEB occurred at a younger age and had a lower mutational burden compared to tumors from patients without RDEB (Extended Data Fig. 1d). On multivariate analysis, mutational burden was marginally associated with shared mutations (p = 0.058) and neither age nor RDEB status were associated with shared mutations. Overall, the two most frequently occurring mutations were found in only seven out of the 149 tumors (4.7%), and 74 out of 149 (49.7%) tumors contained at least one of the 27 shared mutations (Fig. 1b). Thus, a therapeutic approach targeting the 27 shared mutations is expected to cover a maximum of half of the patients with cSCC.

**Figure 1:**
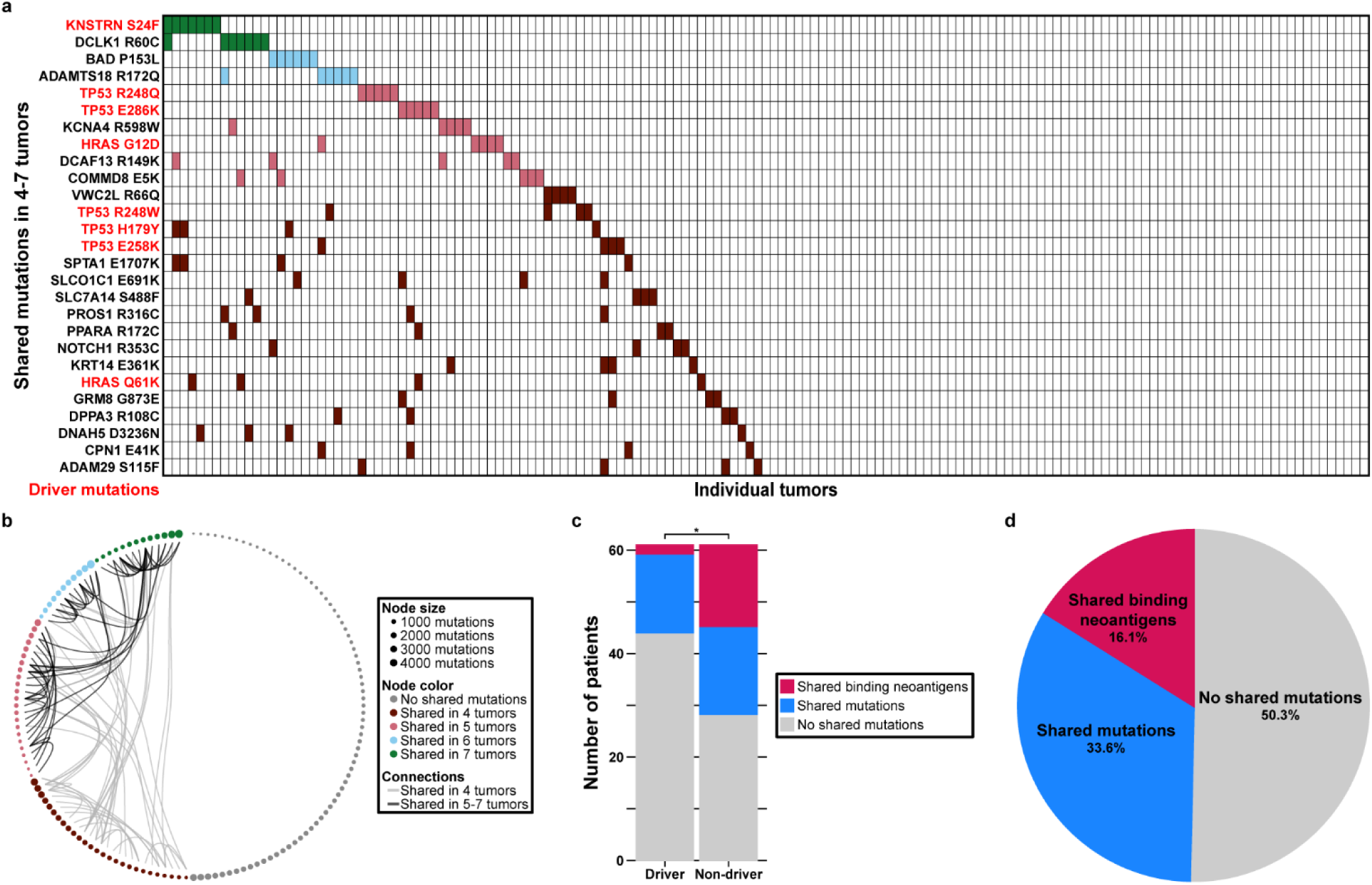
Majority of mutations in human cSCC are unique to an individual tumor. **a,** Waterfall plot of shared missense mutations across 149 human cSCC tumors. Single nucleotide variants were analyzed from two datasets. Shared mutations were defined as mutations that occurred in at least four tumors and at least one tumor from each dataset. Mutations annotated as putative driver mutations are shown in red. **b**, Visualization of the cSCC tumors that contain the 27 shared mutations from (**a**). Each node represents a single tumor. Tumors with a mutation shared in less than four tumors or in only one dataset are annotated as having no shared mutations. **c,** Stacked bar graph displaying the number of patients with putative driver mutations and non-driver mutations that are predicted to result in shared binding neoantigens, shared mutations, and no shared mutations. Number of shared binding neoantigens from driver mutations versus non-driver mutations compared using a chi-squared test. Shared binding neoantigens were defined as missense mutations that were predicted to bind a patient-specific HLA allele with a binding affinity of <500nM. Only tumors for which patient-specific HLA alleles were able to be determined were included. **d,** Pie chart depicting the percentages of patients with predicted shared binding neoantigens, shared mutations, or no shared mutations. Binding affinity for predicted neoantigens was predicted for either the patient-specific HLA alleles, when available, or for the top two most common European HLA-A, HLA-B, and HLA-C alleles. *p < 0.05

While approximately half of the tumors contain a mutation shared with four or more tumors, the potential for these mutations to elicit a T cell response depends on the ability of the resulting peptides to bind MHC class I. Therefore, we predicted the MHC class I binding of each of the shared mutations.^22^ We first predicted binding for the subset of patients for which patient-specific HLA alleles could be determined. Consistent with previous work showing an inverse relationship between oncogenicity and immunogenicity,^22^ the proportion of predicted shared binding neoantigens out of all shared mutations was lower for previously identified driver mutations compared to non-driver mutations (Fig. 1c). To predict the total number of patients with shared binding neoantigens, we predicted the binding for the subset of patients without annotated HLA alleles using the most common European HLA alleles (Extended Data Fig. 1e). European alleles were selected as the prevalence of cSCC is highest in individuals of European descent.^23^ Overall, 24/149 patients (16.1%) contain a shared mutation that is predicted to bind well to MHC class I (Fig. 1d). Furthermore, binding predictions alone likely overestimate the rate of immunogenicity, as previous work has demonstrated that <15% of prioritized neoantigens elicit an immune response, and it is unknown what percentage of these mediate tumor control.^8^

### Mouse cSCC model recapitulates human disease

To address the need for a transplantable, physiologic model to evaluate neoantigens that mediate tumor control, we developed three cSCC cell lines from solar UV light induced invasive cSCC tumors in BALB/c mice (Fig. 2a).^19^ We evaluated the extent to which the murine cSCC cell lines recapitulate human disease through the mutational burden, mutational profile, and driver mutations. The number of total and missense mutations in the murine cSCC cell lines is within the range observed in human cSCC (Fig. 2b). Human tumors and the murine cell lines demonstrate a consistently high UV-induced C>T mutational signature with no significant difference in the percentage of C>T mutations for human cSCC compared to the murine cell lines (Fig. 2b).

**Figure 2:**
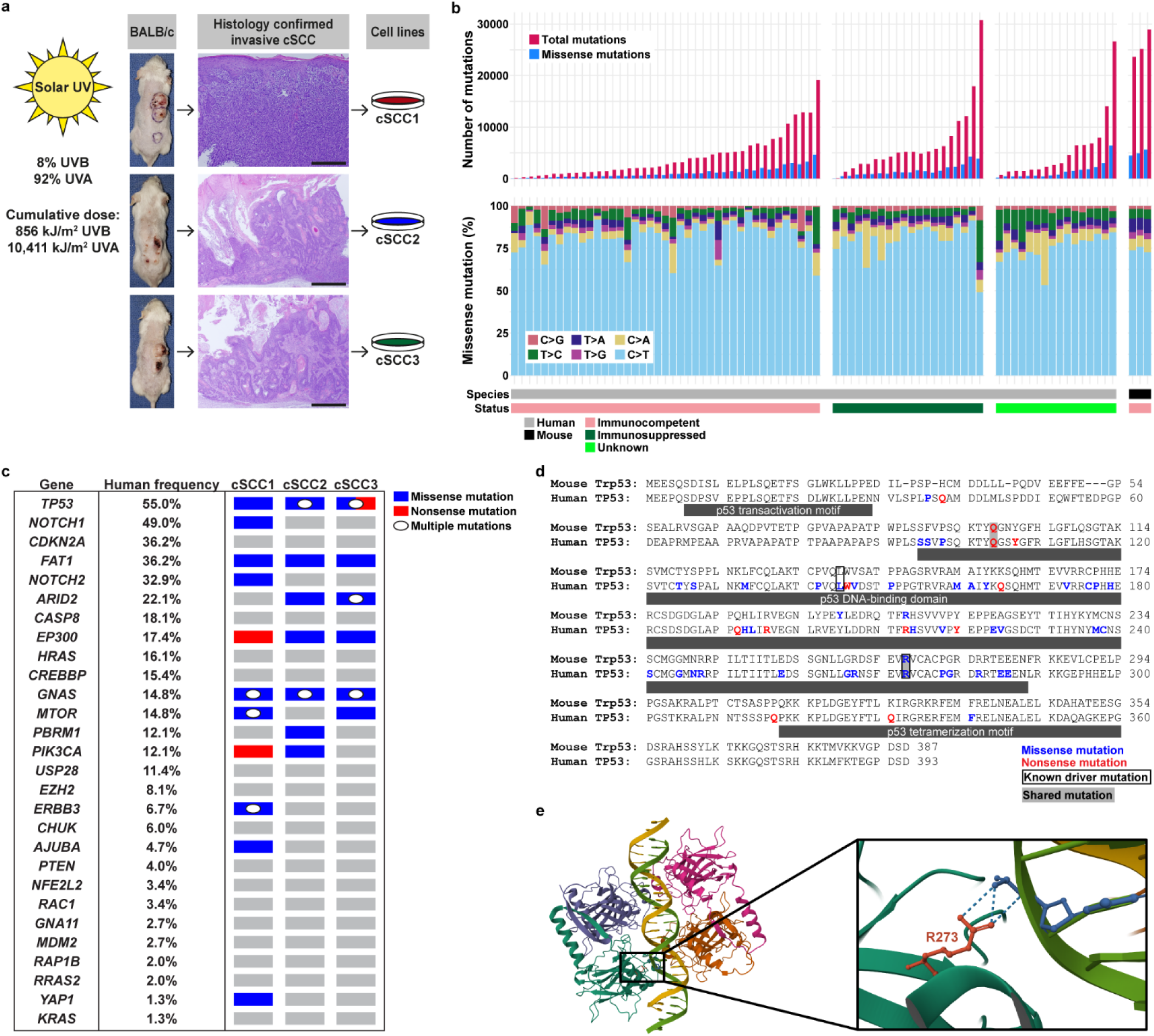
Generation of mouse cSCC model that recapitulates UV mutational signature and driver mutations found in human disease. **a,** BALB/c mice were exposed to solar ultraviolet (UV) light thrice weekly for 42 weeks. Tumors were histologically confirmed as invasive cSCC, and the tumor cells were adapted to culture. **b,** Mutations were identified in 77 human cSCC tumors and the three murine cell lines. Top: Bar graph displaying the total and missense mutations in human and murine cSCC. Bottom: Stacked bar graph displaying the substitution mutations identified in human and murine cSCC. **c,** Comparison of mutated driver genes in human and mouse cSCC. Genes containing driver mutations previously identified in human cSCC are annotated by the frequency of missense or nonsense mutations in human tumors. Right three columns display the presence of missense and/or nonsense mutations in these genes in the three murine cell lines. **d,** Alignment of human *TP53* and mouse *Trp53* showing the location of missense and nonsense mutations. **e,** Visualization of a publicly available crystal structure of the P53 tetramer complexed with DNA. The inset shows the R273 residue that is mutated in one human tumor and two murine cSCC cell lines. Dotted lines indicate non-covalent interactions between the R273 residue and the nucleotide backbone.

Genes that contain known driver mutations in human disease also show frequent mutations in the murine cSCC cell lines (Fig. 2c). In particular, each of the three murine cSCC cell lines contain a *Trp53* mutation (mouse homolog of human *TP53*) that is in the same location as in human disease (Fig. 2d). cSCC3 contains a truncating mutation (Q98* *Trp53*) shared with a human tumor (Q104* *TP53*) that eliminates the binding domain. cSCC1 and cSCC2 contain a missense mutation in the DNA binding domain (R267C *Trp53*) which is identified at the same location in a human tumor (R273H *TP53*). On evaluation of the crystal structure of P53 complexed with DNA, the R273 residue forms non-covalent interactions with the nucleotide backbone of DNA and is important for the function of P53 (Fig. 2e).^24^ Overall, the mouse model recapitulates both the mutational signature and driver mutations of human disease.

### Immunogenic cSCC model constrained dominantly by CD8 T cells with support from CD4 T cells

To study the T cell-mediated response to cSCC, we generated clonal cSCC cell lines (Fig. 3a). The panel of clonal cell lines had high constitutive *in vitro* expression of MHC class I (Extended Data Fig. 2a), and thus these cell lines are subject to CD8 T cell-mediated killing. To assess the extent to which T cells constrain the *in vivo* growth of the panel of clonal cSCC cell lines, we compared the tumor growth between athymic mice, lacking mature T cells, and wild-type mice. Following intradermal injection with each of the six clonal cSCC cell lines, athymic mice had a greater rate of tumor growth and more quickly reached the tumor volume endpoint compared to wild-type mice (Extended Data Fig. 2b-m). These data support that T cells constrain *in vivo* cSCC growth. The panel of cSCC cell lines are highly immunogenic as only a portion of wild-type mice formed tumors with injection of four of the cell lines and no wild-type mice formed tumors with injection of two of the cell lines (Extended Data Fig. 2b-m).

**Figure 3.**
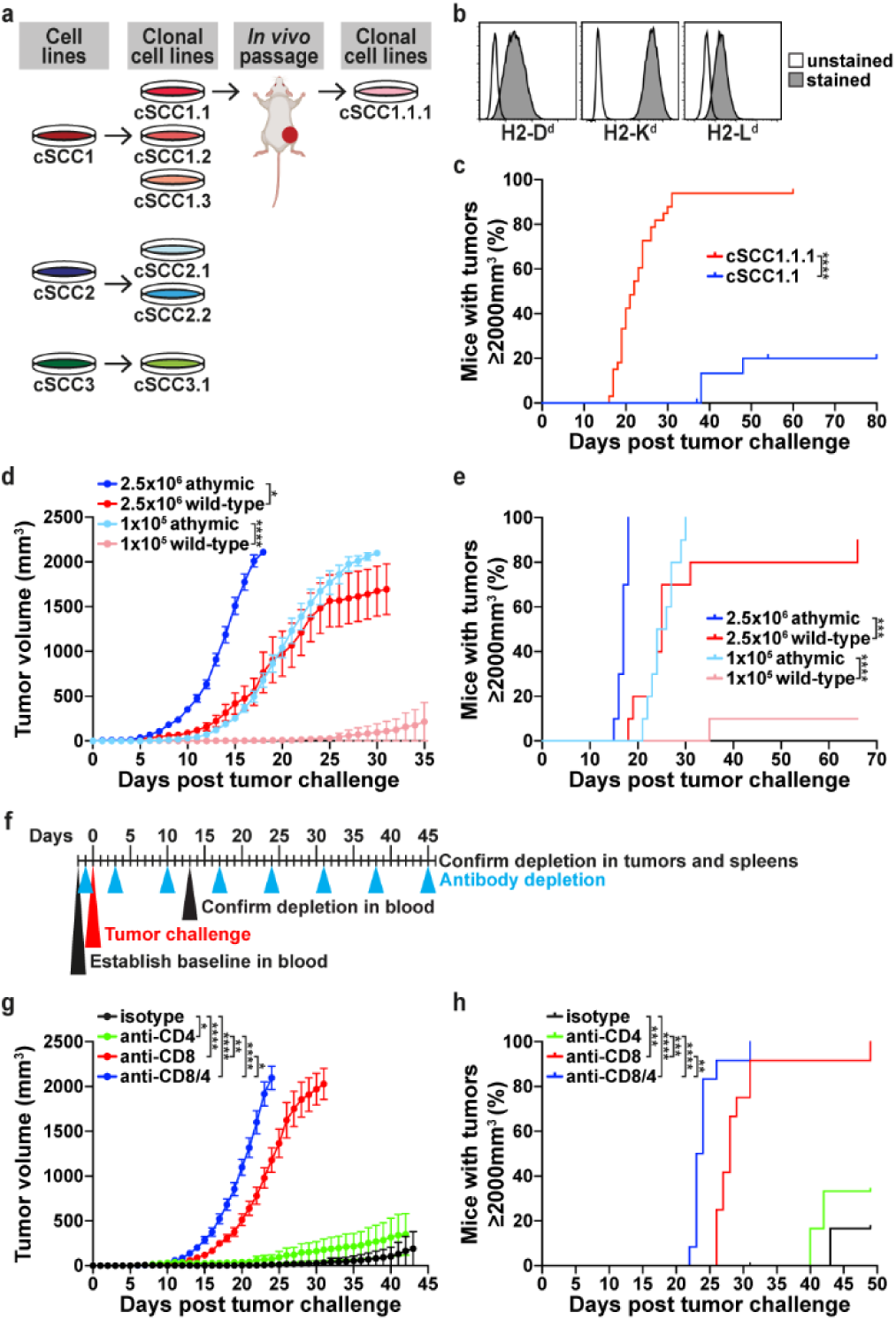
Immunogenic cSCC model constrained by CD8 T cells. **a,** Three cSCC cell lines were used to generate clonal cSCC cell lines. cSCC1.1.1 was generated from an *in vivo* passage of cSCC1.1 in a wild-type mouse, followed by adaptation to culture and cloning. **b,** *In vitro* expression of BALB/c-specific MHC class I alleles in cSCC1.1.1 using flow cytometry. **c,** Wild-type mice were intradermally injected with 2.5××10^6^ cSCC1.1 (n = 20 mice) or cSCC1.1.1 (n = 33 mice) cells. Data pooled from multiple independent experiments. **d,e** Athymic and wild-type mice were intradermally injected with either 1×10^5^ or 2.5×10^6^ cSCC1.1.1 cells (n = 10 per group). Data pooled from two independent experiments. **f,** Schematic of T cell depletion strategy. Mice received intraperitoneal treatment with isotype control, anti-CD4, anti-CD8, or anti-CD8/4 antibodies on days −1 and +3 relative to intradermal injection with 1×10^5^ cSCC1.1.1 cells, and weekly thereafter. **g,h** Tumor volume curves and survival curves for mice (n = 12 per group) described in **f**; data pooled from two independent experiments. Tumor volume (mean ± SEM) over time displayed using last observation carried forward. Tumor growth was compared using a linear mixed effects model and time to reach tumor volume endpoint was compared using a Cox proportional hazards model. *p < 0.05; **p < 0.01; ***p < 0.001; ****p < 0.0001

To create a cell line that more reliably forms tumors *in vivo,* cSCC1.1 was passaged in a wild-type mouse to generate cSCC1.1.1 (Fig. 3a). MHC class I expression was not lost during the *in vivo* passage, as cSCC1.1.1 had high constitutive *in vitro* expression of all three MHC class I alleles (Fig. 3b). cSCC1.1.1 did not express MHC class II with or without IFN-γ treatment (data not shown). Compared to cSCC1.1, cSCC1.1.1 had a faster *in vitro* doubling time, suggesting that the *in vivo* passage resulted in epigenetic changes that enhanced intrinsic growth (Extended Data Fig. 2n). cSCC1.1.1 had faster *in vivo* tumor growth (Extended Data Fig. 2o) and a greater percentage of mice growing tumors (Fig. 3c) compared to cSCC1.1. T cells constrain the *in vivo* tumor growth of cSCC1.1.1, as athymic mice had a greater rate of tumor growth compared to wild-type mice challenged with a low or high number of cSCC1.1.1 cells (Fig. 3d). Additionally, athymic mice reached the tumor volume endpoint sooner than wild-type mice challenged with a low or high number of cSCC1.1.1 cells (Fig. 3e). These data support that cSCC1.1.1 reliably forms tumors *in vivo* and can be used to investigate the T cell-mediated responses that control cSCC.

Flow cytometry of cSCC1.1.1 tumors demonstrated that CD4 T regulatory (Treg) cells are the most numerous intratumoral T cell population; within the tumors there was a 3.4:1 ratio of CD4 to CD8 T cells, a 2.7:1 ratio of CD4 Treg cells to CD4 T conventional cells, and a 2.5:1 ratio of CD4 Treg cells to CD8 T cells (Extended Data Fig. 3). To determine the extent to which CD4 and CD8 T cells constrain the *in vivo* cSCC1.1.1. growth, we treated mice with depleting antibodies prior to tumor challenge with a low cell number (Fig. 3f). We confirmed successful depletion of the targeted T cell population(s) in the blood of each mouse during the experiments (Extended Data Fig. 2p,q). At the end of the experiments or once mice reached the tumor volume endpoint, T cell depletion was confirmed in tumors (data not shown) and/or spleens (Extended Data Fig. 2r,s). Mice depleted of CD8 T cells had a substantial increase in tumor growth compared to mice treated with isotype control (Fig. 3g,h). Depletion of both CD8 and CD4 T cells resulted in a small increase in tumor growth compared with depletion of CD8 T cells alone (Fig. 3g,h). Mice depleted of CD4 T cells had a small but significant increase in tumor growth compared to mice treated with isotype control (Fig. 3g,h). These data support that CD8 T cells play the dominant role and CD4 T cells have a supportive role in constraining cSCC growth.

### Response to vaccination with irradiated cells is tumor specific and dependent on CD8 T cells

We next determined if vaccination with irradiated cSCC cells could constrain tumor growth. To do this, mice were prophylactically vaccinated with irradiated cSCC1.1.1 cells. Mice vaccinated with 5×10^5^ or 2.5×10^6^ cSCC1.1.1 cells irradiated with 10 to 80 Gy had complete protection from tumor challenge with cSCC1.1.1 cells, compared to mice vaccinated with PBS (Extended Data Fig. 4a,b). Additionally, 55 out of 56 mice were protected from a second challenge with cSCC1.1.1 cells 49 days after the initial challenge (data not shown), demonstrating that vaccination resulted in long-lived memory. These results demonstrate that the cSCC1.1.1 cells contain antigens which elicit an adaptive immune response.

To begin to investigate the antigens responsible for the response to vaccination with irradiated tumor cells, we first considered the mutations in the panel of clonal cSCC cell lines. The mutational burden and mutational profile of the clonal cSCC cell lines (Extended Data Fig. 4c) are similar to the cSCC cell lines (Fig. 2b,c). We also found that the majority of the missense mutations (Fig. 4a) and MHC class I binding neoantigens (Fig. 4b) were unique to the three cSCC tumors that were used to generate the clonal cSCC cell lines. Therefore, clonal cSCC cell lines from distinct tumors could be used to interrogate the antigens responsible for response to irradiated tumor cell vaccination. For example, each cSCC cell line likely shares tumor-associated antigens.

**Figure 4:**
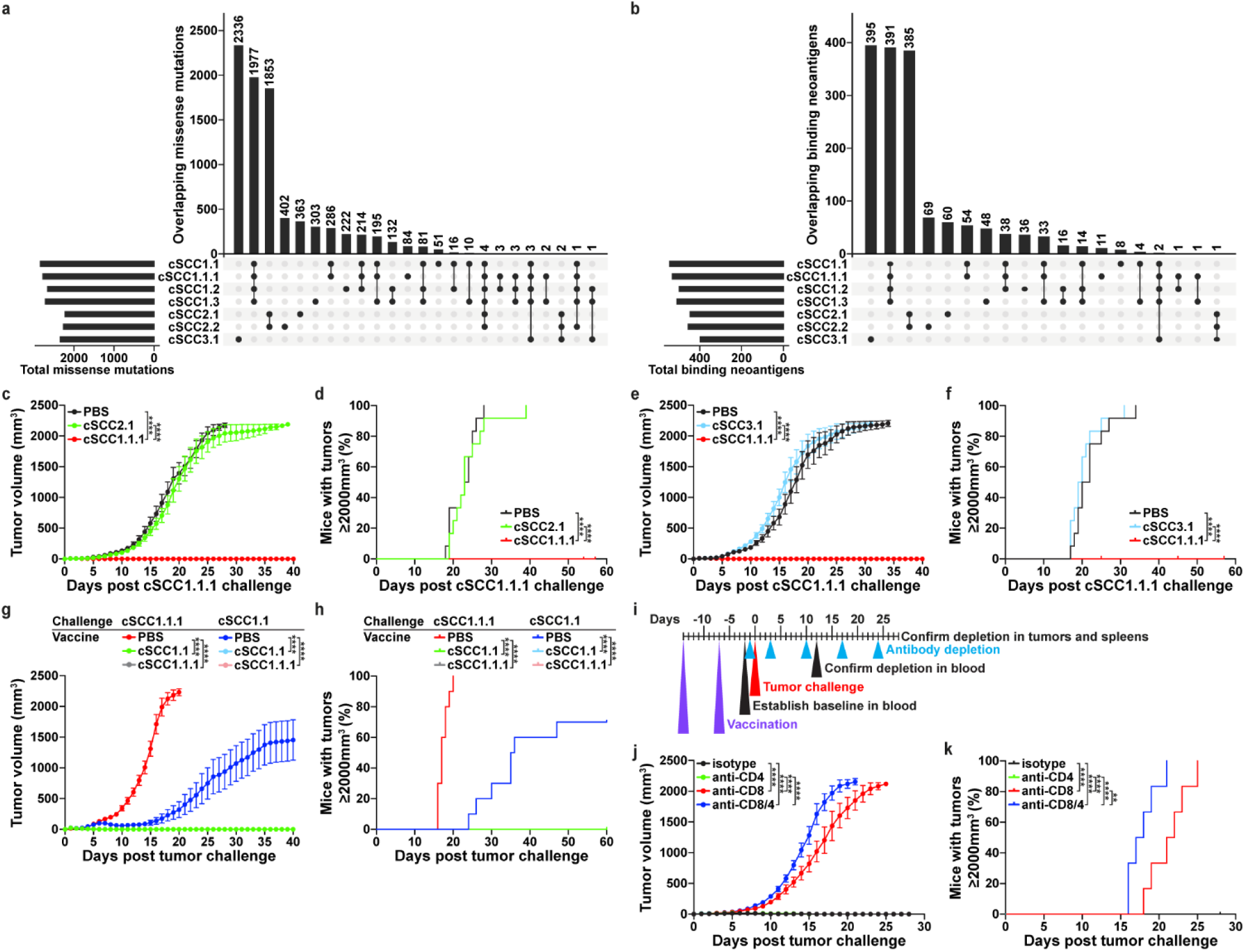
Response to vaccination with irradiated cells is tumor specific and dependent on CD8 T cells. **a,** Upset plot of the missense mutations in the panel of clonal cSCC cell lines. Missense mutations were identified using the overlap of variants called by GATK Mutect2 and Strelka and confirmed in RNA. **b,** Upset plot of the binding neoantigens in the panel of cell lines. Binding neoantigens were defined as mutations with at least one 9mer with a predicted binding affinity rank of <0.5% and eluted ligand rank of <0.5%. **c,** Wild-type mice (n = 12 per group) were intradermally vaccinated with PBS, 5×10^5^ cSCC2.1 cells, or 5×10^5^ cSCC1.1.1 cells irradiated with 80 Gy. One week after vaccination, mice were intradermally injected with 2.5×10^6^ cSCC1.1.1 cells. **d,** Survival curves for mice described in (**c**). **e,** Mice (n = 12 per group) were vaccinated with PBS, 5×10^5^ cSCC3.1 cells, or 5×10^5^ cSCC1.1.1 cells irradiated with 80 Gy and challenged as described in (**c**). **f,** Survival curves for mice described in (**e**). **g,** Mice were vaccinated with PBS, 5×10^5^ cSCC1.1 cells, or 5×10^5^ cSCC1.1.1 cells irradiated with 80 Gy. One week later, mice were challenged with 2.5×10^6^ cSCC1.1.1 cells or 5×10^6^ cSCC1.1 cells. Data shown are from one experiment with n = 10 mice per group. **h,** Survival curves for mice described in (**g**). **i,** Schematic of vaccination and T cell depletion strategy. Mice were vaccinated with irradiated 5×10^5^ cSCC1.1.1 cells on days −14 and −7 relative to tumor challenge with 2.5×10^6^ cSCC1.1.1 cells. Mice received intraperitoneal treatment with isotype control, anti-CD4, anti-CD8, or anti-CD8/4 antibodies on days −1 and +3, and weekly thereafter. **j,** Tumor growth curves and **k,** survival curves for mice described in (**i**). Data shown are from one experiment with n = 6 mice per group. Tumor growth data are pooled from two independent experiments unless stated otherwise. Tumor volume (mean ± SEM) over time displayed using last observation carried forward. Tumor growth was compared using a linear mixed effects model and time to reach tumor volume endpoint was compared using a Cox proportional hazards model. **p < 0.01; ****p < 0.0001

Cell lines cSCC2.1 and cSCC1.1.1 share four missense mutations, including a putative driver mutation *Trp53* R267C (Fig. 4a), but do not share any predicted binding neoantigens (Fig. 4b). Prophylactic vaccination with irradiated cSCC2.1 cells did not protect mice from tumor challenge with cSCC1.1.1 (Fig. 4c,d), supporting that antigens deriving from shared mutations or tumor-associated antigens are not responsible for the response to vaccination with irradiated cSCC1.1.1 cells. To determine if the two predicted binding mutations shared between cSCC1.1.1 and cSCC3.1 (Fig. 4b) contribute to the response to irradiated tumor cell vaccination, mice were prophylactically vaccinated with irradiated cSCC3.1 cells prior to tumor challenge with cSCC1.1.1 (Fig. 4e,f). Vaccination with irradiated cSCC3.1 cells did not provide protection from challenge with cSCC1.1.1, support that the shared predicted binding mutations are not responsible for the protection offered by vaccination with irradiated cSCC1.1.1 cells. Thus, the antigens responsible for the response to vaccination with irradiated cSCC1.1.1 cells are most likely specific to the tumor used to generate cSCC1.1.1.

To further support that the response to vaccination is specific to antigens unique to the tumor that the cells are derived from, mice were vaccinated with irradiated cSCC1.1 or cSCC1.1.1 cells and then challenged with either cSCC1.1 or cSCC1.1.1 (Fig. 4g,h). Vaccination with either clonal cell line completely protected all mice from tumor challenge with either clonal cell line. These data support that the antigens responsible for protection from tumor growth are unique to the tumor. These findings also support that the *in vivo* passage of cSCC1.1 to generate cSCC1.1.1 did not result in the loss of the immunogenic antigens that are responsible for the response to vaccination.

To determine the contribution of CD8 and CD4 T cells to the response to vaccination with irradiated cSCC1.1.1 cells, both the priming and effector phase of the response to vaccination were considered. To evaluate the priming phase, mice were depleted of CD8 and/or CD4 T cells prior to vaccination with irradiated cSCC1.1.1 cells (Extended Data Fig. 4d). Mice depleted of CD8 T cells alone or in combination with CD4 T cells were not protected from tumor challenge by vaccination, supporting that CD8 T cells are responsible for the priming phase of the response to vaccination (Extended Data Fig. 4e-h). The effector phase of the response to vaccination was evaluated by vaccinating mice with irradiated cSCC1.1.1 cells prior to depletion of CD8 and/or CD4 T cells (Fig. 4i). The effector phase of vaccination was dependent on CD8 T cells, as mice depleted of CD8 T cells alone or in combination with CD4 T cells were not protected from tumor challenge by vaccination (Fig. 4j,k; Extended Data Fig. 4i,j). These data support that CD8 T cells have the dominant role in the protective response to irradiated tumor cell vaccination in cSCC. Taken together, these data suggest that protection is mediated by CD8 T cell responses to tumor-specific neoantigens.

### Vaccination with mutant Kars and Picalm.2 constrains *in vivo* cSCC1.1.1 growth

Given the predominant CD8 T cell mediated tumor constraint without vaccination (Fig. 3f-h) and with vaccination (Fig. 4i-k; Extended Data Fig. 4d-f), we focused on identifying the MHC class I neoantigens in cSCC1.1.1. In cSCC1.1.1, 5,010 missense mutations were identified (Fig. 5a). Of these, 2,341 mutations were confirmed in the transcriptome. Since prior work from our group^16^ and others^9, 14, 15^ has shown the importance of the neoantigen-MHC interaction and expression in the prediction of immunogenic neoantigens, we prioritized mutations with high MHC binding affinity (binding affinity % rank <0.5%),^25, 26^ MHC presentation (eluted ligand % rank <0.5%),^25, 26^ and expression (top 5% of variant allele-specific expression). These criteria resulted in 28 prioritized mutations, which were validated via Sanger sequencing and resulted in 30 predicted immunogenic 9mer MHC class I neoantigens.

**Figure 5:**
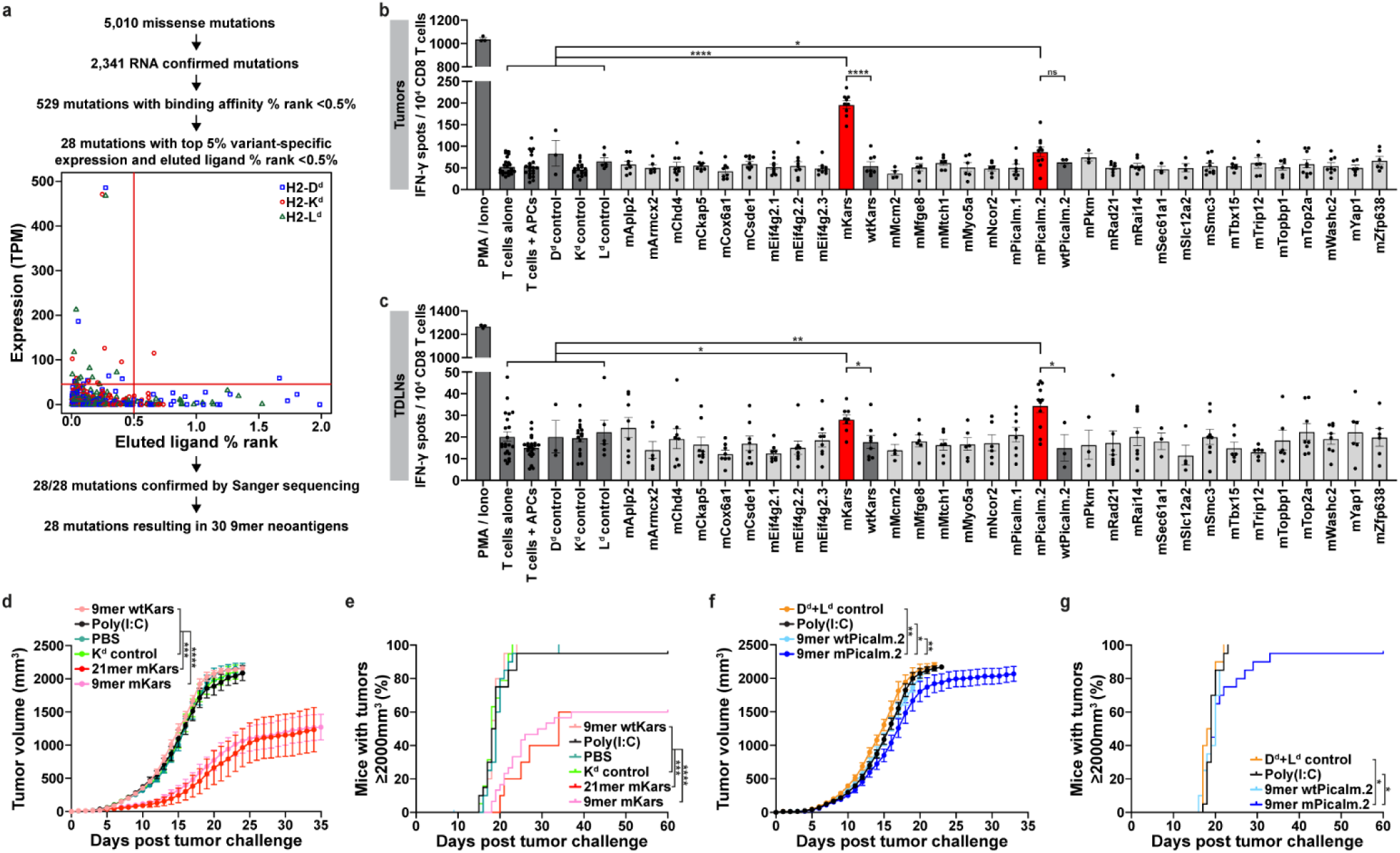
Vaccination with mutant Kars and Picalm.2 constrains *in vivo* cSCC1.1.1 growth. **a,** Workflow to prioritize predicted immunogenic MHC class I neoantigens in cSCC1.1.1. Prioritized immunogenic neoantigens are located in the upper left quadrant, and symbols represent the top neoantigen candidate for a given missense mutation. **b,c** IFN-γ enzyme-linked immune absorbent spot (ELISPOT) results for CD8 T cells isolated from (**b**) tumors and (**c**) tumor-draining lymph nodes (TDLNs) harvested ten days after wild-type mice were challenged with 2.5×10^6^ cSCC1.1.1 cells. CD8 T cells were co-cultured with naïve splenocytes (APCs) pulsed with 2 µg/mL of the indicated individual peptides. In cases where there were multiple mutations in the same protein or the same mutation was in a different position in the peptide, peptides were sequentially named.1, .2, etc. Data displayed as mean ± SEM (n = 3 - 9 independent experiments). IFN-γ spots for all negative control groups (T cells alone, T cells + APCs, and irrelevant binding peptides: D^d^ control, K^d^ control, and L^d^ control) were pooled and compared to each neoantigen peptide using a Kruskal-Wallis test. Mutant (m) peptides were compared to wild-type (wt) peptides using a Mann-Whitney test. Peptides with significantly increased number of IFN-γ spots displayed in red. Controls are displayed in dark grey. **d,** Tumor volume curves for cSCC1.1.1-tumor-bearing mice prophylactically vaccinated with 9mer wtKars with poly(I:C), poly(I:C) alone, PBS, K^d^ control with poly(I:C), 21mer mKars with poly(I:C), or 9mer mKars with poly(I:C) (n = 10 - 20 mice per group). **e,** Survival curves for mice described in (**d**). **f,** Tumor volume curves for cSCC1.1.1-tumor-bearing mice prophylactically vaccinated with D^d^ and L^d^ control with poly(I:C), poly(I:C) alone, 9mer wtPicalm.2 with poly(I:C), or 9mer mPicalm.2 with poly(I:C) (n = 10 - 20 mice per group). **g,** Survival curves for mice described in (**f**). Mice in prophylactic vaccination experiments received 100 µg poly(I:C) ± 50 µg peptide on days −14 and −7 relative to tumor challenge with 2.5×10^6^ cSCC1.1.1 cells. Tumor volume (mean ± SEM) over time displayed using last observation carried forward. Tumor growth was compared using a linear mixed effects model. Time to reach tumor volume endpoint was compared using a Cox proportional hazards model. ns (not significant); *p < 0.05; **p < 0.01; ***p < 0.001; ****p < 0.0001

Of the 30 predicted immunogenic MHC class I neoantigens, two neoantigens elicited IFN-γ+ CD8 T cell responses by enzyme-linked immune absorbent spot (ELISPOT) analysis. Both mKars and mPicalm.2 induced more IFN-γ+ producing CD8 T cells derived from cSCC1.1.1 tumors (Fig. 5b) and tumor-draining lymph nodes (TDLNs) (Fig. 5c) compared to T cells alone; T cells and antigen presenting cells (APCs); and irrelevant binding peptides: D^d^ control, K^d^ control, and L^d^ control. To assess whether Kars-specific CD8 T cells and Picalm.2-specific CD8 T cells selectively recognize the mutant peptide sequences, the mutant and wild-type peptides were compared in ELISPOT. Compared to wtKars, mKars elicited more IFN-γ+ CD8 T cells in both tumors and TDLNs; and, mPicalm.2 induced more IFN-γ+ CD8 T cells in TDLNs compared to wtPicalm.2. The IFN-γ+ CD8 T cell responses to mKars and mPicalm.2 were not observed in CD8 T cells isolated from naïve lymph nodes (LNs) (Extended Data Fig. 5a). Additionally, none of the neoantigens elicited IFN-γ+ CD4 T cell responses in tumors, TDLNs, or naïve LNs (Extended Data Fig. 5b-d). This supports that both mKars and mPicalm.2 are immunogenic neoantigens.

As mKars and mPicalm.2 elicited IFN-γ+ CD8 T cell responses, we tested whether vaccination with mKars or mPicalm.2 could protect against cSCC1.1.1 tumor outgrowth. Mice prophylactically vaccinated with either 9mer or 21mer mKars with adjuvant poly(I:C) had a slower rate of tumor growth compared to mice vaccinated with wtKars, poly(I:C) alone, PBS, or K^d^ control (Fig. 5d). Vaccination with 9mer mKars or 21mer mKars protected 40% of mice against tumor outgrowth (Fig. 5e). Although some murine studies combine peptide vaccines with immune checkpoint inhibitors,^17, 27^ we demonstrate that these tumor-rejecting neoantigens constrain tumor growth in the absence of immune checkpoint inhibitors. The similar response to vaccination with 9mer or 21mer mKars supports that the 9mer mKars peptide represents the immunogenic T cell epitope and that the response is primarily mediated by CD8 T cells. Progressively growing tumors in mice vaccinated with mKars were evaluated for mechanisms that may have contributed to tumor escape. The heterozygous Kars mutation was retained in all tumors evaluated from mKars and poly(I:C) vaccinated mice (data not shown). Similarly, Kars mRNA expression was not significantly different in tumors that escaped mKars vaccination compared to tumors from mice vaccinated with poly(I:C) (Extended Data Fig. 6a). Finally, there was no significant decrease in the expression of H2-K^d^, the predicted binding allele for mutant Kars, on tumor cells between poly(I:C) and mKars vaccinated tumors (Extended Data Fig. 6b). These data show that tumor escape was not due to genomic loss of mKars DNA, loss of *Kars* RNA expression or loss of MHC class I; tumor escape may be due to other immunosuppressive changes in the tumor microenvironment. Prophylactic vaccination with 9mer mPicalm.2 together with poly(I:C) resulted in delayed tumor growth compared to vaccination with D^d^ and L^d^ control, poly(I:C) alone, or wtPicalm.2 (Fig. 5f). Vaccination with mPicalm.2 resulted in a small but significant delay in mice reaching the tumor volume endpoint; however, only one mouse had complete protection from tumor outgrowth (Fig. 5g). Vaccination with wtKars or wtPicalm.2 did not constrain tumor growth or protect mice from tumor challenge, demonstrating that the response to neoantigen vaccination was specific to the mutations. These data demonstrate that mKars and mPicalm.2 are tumor-rejecting neoantigens, and that mKars is a superior tumor-rejecting neoantigen.

### Structural differences relative to wild-type and improved MHC binding are features of tumor-rejecting neoantigens

Next, we evaluated the interactions of the tumor-rejecting neoantigens, mKars and mPicalm.2, with MHC class I alleles to gain insights into the mechanisms underlying the ability to mediate tumor rejection. We first experimentally determined the ability of the tumor-rejecting neoantigens to stabilize MHC class I alleles on the cell surface of transporter associated with antigen processing (TAP)-deficient T2 cells. Both mKars and wtKars strongly stabilized H2-K^d^ (Fig. 6a), but not H2-L^d^ (Fig. 6b) or H2-D^d^ (Fig. 6c). The mKars, wtKars, and K^d^ binding control peptides had similar binding affinities for H2-K^d^ (Fig. 6d). Compared to the L^d^ binding control peptide, mPicalm.2 strongly stabilized H2-L^d^, whereas wtPicalm.2 only moderately stabilized H2-L^d^ (Fig. 6b). The relative binding affinity of mPicalm.2 to H2-L^d^ was weaker than the L^d^ binding control, but stronger than wtPicalm.2 (Fig. 6e). Both mPicalm.2 and wtPicalm.2 similarly stabilized H2-D^d^ (Fig. 6c) and had a decreased relative binding affinity to H2-D^d^ compared to the D^d^ binding control (Fig. 6f). Of note, the experimentally observed levels of binding of mPicalm.2 and wtPicalm.2 to H2-D^d^ was not predicted (Fig. 6g). Together these data support that 1) mKars and wtKars peptides equivalently bind H2-K^d^, 2) mPicalm.2 peptide binds H2-L^d^ better than wtPicalm.2, and 3) unexpectedly, mPicalm.2 and wtPicalm.2 similarly bind H2-D^d^.

**Figure 6:**
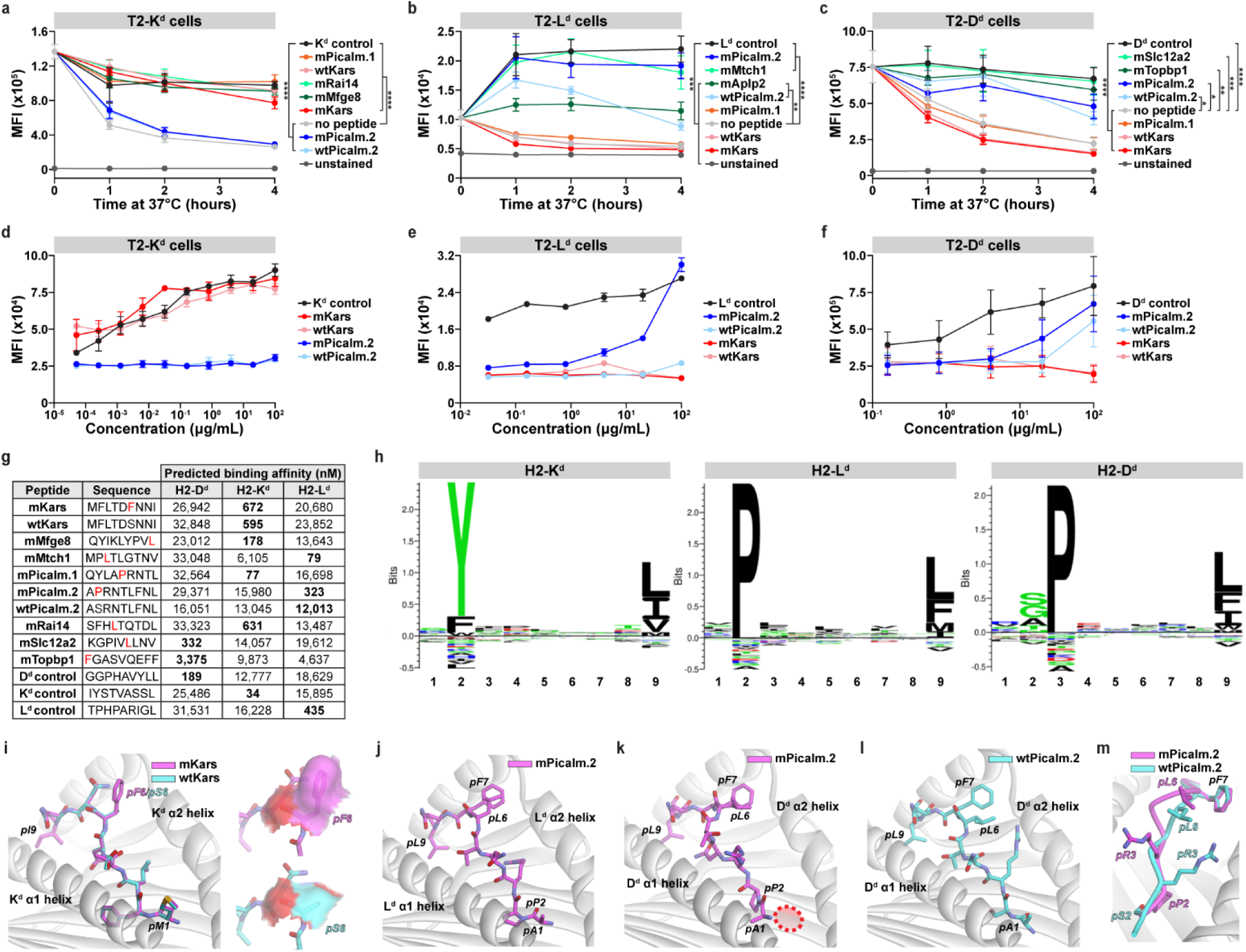
Structural differences relative to wild-type and improved MHC binding are features of tumor-rejecting neoantigens. **a-c,** Median fluorescence intensity (MFI) of MHC class I allele staining measured via flow cytometry for (**a)** T2-K^d^ cells, (**b**) T2-L^d^ cells, and (**c**) T2-D^d^ cells pulsed with 100 μg/mL of the indicated peptides over time. Data displayed as mean ± SEM (n = 3 independent experiments). **d-f**, MFI of MHC class I staining for (**d**) T2-K^d^ cells, (**e**) T2-L^d^ cells, and (**f**) T2-D^d^ cells pulsed with various concentrations of the indicated peptides. Data displayed as mean ± SEM (n = 3 independent experiments) and analyzed using linear regression. **g,** Table with NetMHCpan predicted binding affinities for mutant (m) and wild-type (wt) peptides used in MHC class I stability and binding experiments. Red amino acids show the mutation. The best predicted binding allele is bolded. **h,** NetMHCpan eluted ligand motifs for naturally presented ligands on BALB/c-specific MHC class I alleles, which show the preferred amino acid at each position. **i-m** Structures of the peptide:MHC complexes generated via AI-based modeling. (**i**) mKars and wtKars presented by H2-K^d^. Left panel shows the models of the two complexes superimposed by the H2-K^d^ peptide binding groove, revealing their identical conformations in the groove. The right panel shows the solvent accessible surface areas of the position 6 (p6) amino acid, illustrating the greater exposed aromatic surface for mKars (top) compared to wtKars (bottom). (**j**) mPicalm.2 presented by H2-L^d^ with the aromatic and hydrophobic amino acids at p6 and p7 fully exposed; (**k**) mPicalm.2 presented by H2-D^d^. The model shows the proline at p2 occupying the C pocket, with the A pocket empty as illustrated by the dashed oval, leading to a register shifted conformation. P6 and p7 are again fully exposed; (**l**) wtPicalm.2 presented by H2-D^d^, with the A pocket occupied by the p1 alanine; and (**m**) Complexes of mPicalm.2 and wtPicalm.2 bound to H2-D^d^ superimposed by the peptide binding groove, highlighting the conformational differences between mutant and wild-type resulting from the mutant register shift. *p < 0.05; **p < 0.01; ***p < 0.001; ****p < 0.0001

To examine predicted interactions of the tumor-rejecting neoantigens with MHC and the TCR as well as structural correlates with tumor rejection, we modeled the neoantigen conformations when bound to their presenting MHC class I proteins. For modeling, we used an AI-based approach which we previously demonstrated significantly outperformed legacy, physics-based modeling of peptide/MHC complexes.^28, 29^ Since the tumor rejection mediated by mKars is not explained by improved binding to H2-K^d^ compared to wtKars, we used structural modeling to assess possible differences in the conformation of mKars and wtKars bound to H2-K^d^. Both mKars and wtKars share the position two (p2) phenylalanine and p9 isoleucine (Fig. 6g), which serve as primary anchors for H2-K^d^ (Fig. 6h). The model of mKars bound to H2-K^d^ shows a bulged conformation typical of a nonamer (Fig. 6i, left). The mutated p6 phenylalanine extends up and away from the groove, leaving the side chain fully exposed for engagement by TCRs. We also modeled the wtKars peptide to evaluate any structural differences resulting from the mutation of serine to phenylalanine at p6. The wtKars peptide is predicted to adopt the same conformation as the neoantigen, except for the wild-type serine at p6, whose side chain points down, forming a hydrogen bond to the base of the H2-K^d^ α2 helix. The difference in the structure and chemistry of the p6 side chain results in mKars exposing significantly more hydrophobic and aromatic surface than wtKars (Fig. 6i, right). Notably, exposed aromatic side chains are associated with strong TCR binding and peptide immunogenicity.^30, 31, 32^ Thus, the ability of mKars presented on H2-K^d^ to mediate tumor rejection is predicted to be due to improved TCR recognition of mKars compared to wtKars.

We next used modeling to evaluate the interaction of mPicalm.2 bound to H2-L^d^ as well as H2-D^d^, as mPicalm.2 bound (Fig. 6e,f) and stabilized both alleles (Fig. 6b,c). The p2 proline in mPicalm.2 is the preferred residue at the primary anchor for H2-L^d^, thus explaining the improved binding of mPicalm.2 compared to wtPicalm.2. The H2-L^d^ model of mPicalm.2 showed a typical, bulged nonameric conformation (Fig. 6j), with the p2 proline and p9 leucine occupying the H2-L^d^ B and F pockets, respectively. The side chains of the p6 leucine and p7 phenylalanine are predicted to extend up from the peptide bulge, again displaying significant aromatic and hydrophobic surface that are often associated with TCR binding and immunogenicity (Fig. 6j).^31, 32^ Together these data support that the ability of mPicalm.2 presented by H2-L^d^ to mediate tumor rejection is most likely due to improved binding affinity for H2-L^d^ compared to wtPicalm.2.

The models were particularly helpful in understanding the binding (Fig. 6c) and stabilization (Fig. 6f) of H2-D^d^ by both mPicalm.2 and wtPicalm.2, which was not predicted by NetMHCpan (Fig. 6g). The model of mPicalm.2 bound to H2-D^d^ revealed that rather than adopting a traditional nonameric binding geometry, the peptide was predicted to register shift, or adopt a decameric conformation, resulting in the mutant p2 proline occupying the H2-D^d^ C pocket and the p1 alanine in the B pocket (Fig. 6k). This leaves the A pocket empty but allows the peptide to match the H2-D^d^ peptide binding motif, which shows a strong preference for a proline for the C pocket, usually found in peptide p3 (Fig. 6h), but instead found at p2 in mPicalm.2. We also modeled wtPicalm.2 to H2-D^d^. The wtPicalm.2 peptide adopted a nonameric binding geometry as expected, with the p1 alanine in the A pocket and the p2 serine in the B pocket (Fig. 6l). The register shift in mPicalm.2 leads to substantial conformational differences in the peptides from the N-terminus through to the p7 phenylalanine, resulting in the presentation of a significantly altered peptide configuration from p3 to p6 in mPicalm.2 compared to wtPicalm.2 (Fig. 6m). The register shifted conformation for mPicalm.2 readily explained its binding to H2-D^d^ despite its classification as a nonbinder by NetMHCpan. The similar levels of binding to H2-D^d^ by wtPicalm.2 and mPicalm.2 are also explained by the models: the mutant leaves the A pocket empty but places a highly preferred proline in the C pocket, whereas wtPicalm.2 occupies the A pocket, but with a suboptimal, upward extending arginine at the C pocket as opposed to the more highly preferred proline. Thus, the ability of mPicalm.2 presented by H2-D^d^ to mediate tumor rejection is likely due TCR recognition of a distinct peptide conformation that differentiates it from wtPicalm.2.

### Tumor-rejecting neoantigens with a low differential agretopic index have increased solvent accessible surface area of the mutated residue compared to the corresponding wild-type residue

We next sought to determine the extent to which the structural features and the differences from wild-type for mKars bound to H2-K^d^ and mPicalm.2 bound to H2-D^d^ are generalizable characteristics of tumor-rejecting neoantigens that are not predicted to alter MHC binding. Structural modeling was performed on all non-immunogenic neoantigens from this study and tumor-rejecting neoantigens from this and existing mouse models.^9, 11, 12, 17, 27^ The dataset of tumor-rejecting neoantigens included neoantigens identified using different prioritization methods across different mouse MHC alleles (H2-K^d^, H2-L^d^, H2-D^d^, H2-K^b^, H2-D^b^, and H2-K^k^) and cancer models (cSCC, head and neck SCC, colon cancer, and three chemically-induced sarcomas). The solvent accessible surface area (SASA) was determined for the peptides as well as for the mutated residue and corresponding wild-type residue for each neoantigen. We split the neoantigens by the predicted differential agretopic index (DAI) into a low DAI and high DAI category. The difference in the mutant and wild-type SASA was compared between tumor-rejecting and non-immunogenic neoantigens in both categories. A high DAI means that the neoantigen is predicted to bind MHC better than the wild-type peptide. Since no previous work has defined a criterion for “high DAI,” we searched for a cut point that optimized the p-value for the comparison of the difference in the mutant and wild-type SASA in the low DAI category (Extended Data Fig. 7). The p-value was consistently low across DAI cut points from 1.05 to 1.7 and was optimal at 1.3. Tumor-rejecting neoantigens with a low DAI had a higher difference in SASA between the mutated and wild-type residue than the three other groups (Fig. 7); significance in this analysis was not achieved when either hydrophobic SASA or the SASA of the entire peptide was used, which we believe may stem from the limited size of the database and uncertainties in overall models, particularly for peptides greater than nine amino acids). While a change in the SASA of the mutated residue distinguished the tumor-rejecting neoantigens from the non-immunogenic neoantigens in the low DAI category, there was no difference in the SASA between tumor-rejecting and non-immunogenic neoantigens in the high DAI category. These findings support that the potential for a neoantigen to mediate tumor rejection may result from improved MHC binding or a change peptide structural features that would enhance TCR recognition; the latter is reflected by increased SASA of the mutated residue, as illustrated in the comparison of the mutant and wild-type Kars peptide in Fig. 6i. Overall, this analysis supports the use of structural modeling and SASA as an additional characteristic in prioritizing candidate tumor-rejecting neoantigens in the absence of large changes in MHC binding.

**Figure 7:**
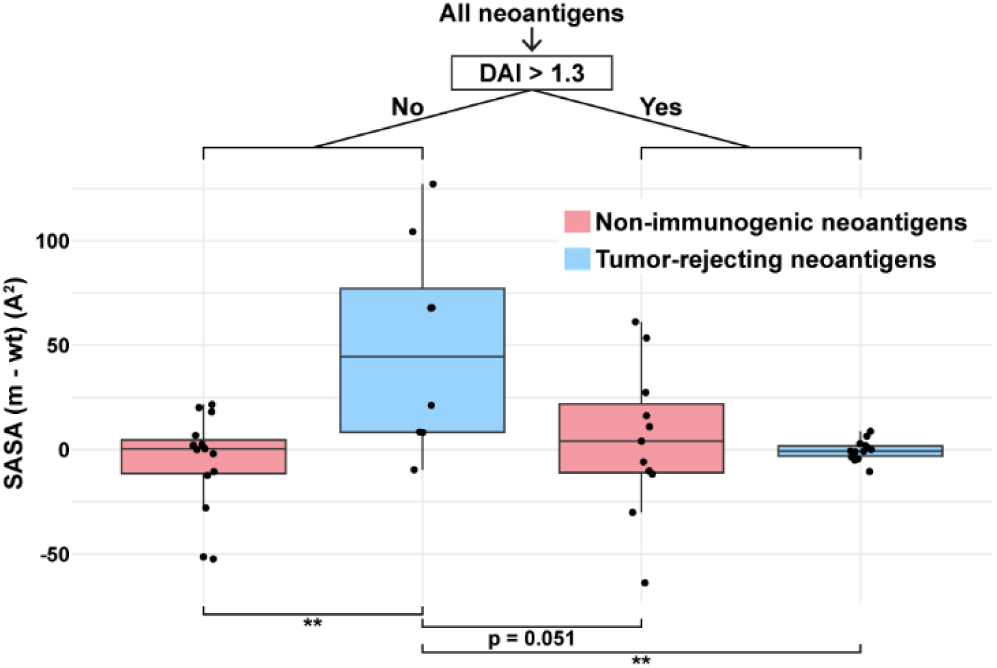
Tumor-rejecting neoantigens with a low differential agretopic index have increased solvent accessible surface area of the mutated residue compared to the corresponding wild-type residue. Non-immunogenic neoantigens (neoantigens with a negative ELISPOT result from this study) were analyzed in combination with tumor-rejecting neoantigens from this and prior studies. Tumor-rejecting neoantigens from prior studies were included if the precise epitope had been individually tested for the ability to elicit tumor-destruction. Neoantigens were split into groups based on their differential agretopic index (DAI), which was calculated as the ratio of the predicted wild-type and mutant peptide dissociation constants. The difference between the solvent accessible surface area (SASA) of the mutant (m) and corresponding wild-type (wt) residues were compared between tumor-rejecting and non-immunogenic neoantigens in both categories using a Wilcoxon Rank Sum test. The DAI split of 1.3 was found to optimize the observed difference in the mutant and wild-type SASA between tumor-rejecting and non-immunogenic neoantigens with low DAI. The bold line indicates the median, and the upper and lower limits of the boxes indicate the 75th and 25th percentiles, respectively. The lower and upper whiskers indicate the minimum and maximum. Dots outside of the box and whiskers indicate outliers. **p < 0.01.

## Discussion

Human cSCC tumors are constrained by T cells, as a portion of cSCC patients respond to immune checkpoint inhibitors,^33^ and immunosuppressed solid organ transplant recipients have a 65-250 fold greater risk of developing cSCC.^34^ However, the lack of response to immune checkpoint inhibition in the majority of patients, the undesirable immune-related adverse events, and the relative contraindication for use of immune checkpoint inhibition in transplant recipients highlights the need for additional immunotherapeutic approaches for cSCC.^33, 34, 35^ Recent work in other cancers has demonstrated the potential for shared neoantigen vaccines.^36, 37^ However, in human cSCC, the vast majority of mutations are unique to each tumor, with the most common mutation shared in only 4.7% of tumors. A theoretical neoantigen vaccine targeting shared neoantigens predicted to bind MHC with a dissociation constant <500nM would cover less than 15% of patients. As only a small percentage of neoantigens predicted to bind MHC stimulate T cell responses,^8, 9, 10, 11, 12, 13, 14, 15, 16^ and an even smaller percentage of neoantigens mediate tumor destruction,^9, 10, 11, 12, 17^ it is likely that significantly less than 15% of patients would demonstrate an appreciable response to a shared neoantigen vaccine in cSCC. The low rate of shared neoantigens and scarcity of immunogenic neoantigens support the need for personalized immunotherapeutic approaches for the treatment of and prevention of recurrence in advanced cSCC.

For personalized neoantigen-based immunotherapies to be effective in cSCC and other high mutational burden tumors, the tumor-rejecting neoantigens need to be identified and targeted. To study tumor-rejecting neoantigens, we generated a transplantable solar UV-induced cSCC model. This cSCC model was generated from tumors induced using solar UV light, which has a similar UV spectral output to sunlight, the etiologic factor that causes human disease.^38^ The cSCC model has a high mutational burden, a high proportion of UV-signature mutations, and shares driver mutations with human disease. Consistent with human disease, the cSCC model is immunogenic as the cSCC model was constrained by T cells and irradiated tumor vaccination completely protected all mice from tumor challenge. Thus, our cSCC model recapitulates human cSCC and can be used to identify characteristics of tumor-rejecting neoantigens that can be applied to human cSCC or other high mutational burden tumors.

We found that CD8 T cells had the dominant role and CD4 T cells had a supporting role in constraining cSCC growth. We also observed that both the primary and effector phases of the response to vaccination with irradiated tumor cells was predominantly driven by CD8 T cells. In tumors from our mouse model, CD4 T cells outnumbered CD8 T cells, and Treg cells outnumbered CD8 T cells, similar to human cSCC.^39^ The fact that intratumoral CD4 Treg cells were more abundant than CD4 T conventional cells may explain why this mouse model was only marginally constrained by CD4 T cells. Consistent with our findings, studies in other cSCC models found a dominant role of CD8 T cells and either no role or a supporting role for CD4 T cells.^40, 41, 42, 43^ In other tumor models, CD4 T cells and MHC class II neoantigens are required for optimal responses to neoantigen-based immunotherapy or response to immune checkpoint inhibitors.^12, 14^ A key difference in these models is that CD4 T cells contribute significantly to the anti-tumor immune responses, as depletion of CD4 T cells alone dramatically increases tumor growth.^12, 14^ Whereas, in our cSCC model, CD4 T cells alone did not dramatically increase tumor growth in the absence of treatment or contribute to the response to vaccination with irradiated tumor cells. These results support that CD8 T cells have the primary role in controlling cSCC in this model, and thus, we focused on the identification of MHC class I neoantigens. Overall, these results suggest that CD8 T cells have the primary role in controlling cSCC.

Accurate prediction and targeting of tumor-rejecting neoantigens is critical for optimizing the efficacy of neoantigen-based immunotherapies for cancer. Recent clinical studies with personalized mRNA cancer vaccines demonstrate the feasibility of the implementation of patient-specific vaccines and clinical benefit.^2, 3, 6^ However, the precise tumor-rejecting neoantigens that are responsible for improved outcomes observed in a portion of patients is unknown, as pools of neoantigens and long peptides/mRNA have been targeted.^2, 3, 4, 6^ Thus, discrete tumor-rejecting neoepitopes have not yet been identified in humans. Furthermore, most studies evaluating neoantigen immunogenicity and datasets used to develop neoantigen prediction tools define immunogenic neoantigens based on ex vivo methods such as ELISPOT, ELISA, and multimer staining.^4, 8, 15, 44, 45^ These ex vivo methods do not take into account the complex in vivo interactions in the tumor microenvironment needed for T cell-mediated control, and immunogenicity demonstrated by ex vivo methods does not necessarily correspond to tumor control.^11^ Additionally, only a small number of discrete tumor-rejecting neoepitopes have been identified in mouse cancer models.^9, 11, 12, 17, 27^ Here, we identify two additional tumor-rejecting neoantigens to assist in the determination of neoantigen characteristics associated with tumor-rejection.

Prior studies have identified improved MHC binding of the neoantigen relative to the wild-type peptide as a characteristic of tumor-rejecting neoantigens,^11, 27^ leading to more effective antigen presentation and increasing the likelihood of T cell recognition. Consistent with prior work, we found that the tumor-rejecting neoantigen, mPicalm.2, better stabilized H2-L^d^ and had a stronger relative binding affinity compared to wtPicalm.2, resulting in a high DAI. Neoantigens that have a high DAI, such as mPicalm.2 presented on H2-L^d^, typically have a mutation in an anchor residue. Capietto et al. extended these findings by demonstrating that the relative binding affinity only distinguishes immunogenic from non-immunogenic neoantigens in the subset of neoantigens with mutations in anchor residues.^46^ Similarly, in the tumor-rejecting neoantigens evaluated in Fig. 7, we found that tumor-rejecting neoantigens with a mutation in an anchor residue had a significantly higher DAI than tumor-rejecting neoantigens with a mutation in a non-anchor residue (data not shown). While a high DAI was characteristic of the tumor-rejecting neoantigens with a mutation in an anchor residue evaluated in this study, we did not observe that DAI distinguished tumor-rejecting from non-immunogenic neoantigens (data not shown). This result is likely due to the fact that each validated, non-immunogenic neoantigen has high predicted MHC binding affinity, and our dataset of non-immunogenic neoantigens with a mutation in the anchor residue only includes one neoantigen which binds MHC worse than the wild-type peptide. Together, these studies support that, for neoantigens that have a mutation in an anchor residue, improved binding of the mutant peptide compared to the wild-type peptide is an important characteristic of tumor-rejecting neoantigens.

For neoantigens where mutations are not in an anchor residue and do not alter MHC binding, distinction from wild-type is required to overcome T cell tolerance. Several studies have indicated how this can be achieved by mutations that alter peptide exposed surface area or conformation in the groove, structurally differentiating the neoantigen from the wild-type peptide and facilitating TCR recognition.^17, 27, 29, 47^ This is exemplified by mKars; we found that although both mKars and wtKars stabilized and had similar binding affinities to H2-K^d^, the nonanchor mutation in mKars resulted in a large increase in solvent accessible surface area. Prior studies of T cell epitopes derived from self and pathogens (primarily viral, but also some neoantigens) emphasized that aromatic and hydrophobic residues, particularly in residues not related to MHC binding, are associated with improved T cell recognition and immunogenicity.^29, 30, 31, 32, 48^ Consistent with these findings, the mutation in mKars changes an exposed serine to a phenylalanine. However, aromaticity or hydrophobicity did not discriminate between the tumor-rejecting neoantigens and non-immunogenic neoantigens in this study. This outcome may be due to the limited dataset size or uncertainties associated with structural modeling.

Furthermore, in a human dataset limited to neoantigens, hydrophobicity did not distinguish immunogenic and non-immunogenic neoantigens; however, in this study the hydrophobicity limited to TCR contact residues was not specifically assessed.^16^ Nonetheless, our analysis clearly identifies structural differences as reflected by changes in solvent accessibility as a means to differentiate neoantigens from their wildtype counterparts and promote T cell recognition. This is further exemplified by the prediction of a register shift resulting from the mutation in mPicalm.2 when bound to H2-D^d^, whereby the mutant results in the peptide leaving the MHC A pocket empty and adopting a decameric bulged conformation, but wild-type occupies the MHC A pocket and adopts a traditional nonameric conformation. We observed a similar register shift in human HLA-A2, where a nonameric tumor antigen with a leucine at p1 adopted a decameric conformation, with the p1 leucine occupying the HLA-A2 B pocket and leaving the A pocket empty and leading to a substantially different peptide conformation and surface exposure,^49^ as predicted for mutant and wild-type Picalm.2 binding H2-D^d^ here. As a general means to capture structural changes from wild-type due to mutation, we tested the extent to which solvent accessibility of TCR contact residues is a predictor of tumor-rejection and found that an increase in the solvent accessibility of the mutated residue relative to the wild-type residue distinguished tumor-rejecting from non-immunogenic neoantigens across mouse cancer models, MHC alleles and neoantigen prioritization strategies.

Improving the clinical efficacy of personalized neoantigen-based immunotherapies requires identification of neoantigens that mediate tumor rejection. This work characterized the neoantigen landscape in human cSCC, presents a novel, clinically-relevant cSCC mouse model and extends the limited number of validated precise neoepitopes with the ability to mediate tumor rejection. This is the first study, to our knowledge, which has evaluated characteristics generalizable to all known murine tumor-rejecting neoantigens. We found that within the category of neoantigens that do not have a major impact on MHC binding, higher solvent accessibility of the mutated amino acid compared to the wild-type counterpart was characteristic of tumor-rejecting neoantigens. While some characteristics, like neoantigen/MHC binding affinity and stability and expression of the neoantigen can be applied to all neoantigens to predict immunogenicity,^15, 16, 26, 46^ this work extends the conceptual framework that there are classes of neoantigens for which different sets of characteristics drive immunogenicity. Namely, improved MHC binding is important for mutations in anchor residues, and improved solvent accessibility is important for mutations in non-anchor residues. Application of these characteristics may improve the selection of neoantigens for inclusion in vaccines for cSCC and other cancers. These findings are most important to apply in high mutational burden cancers, especially those like cSCC that do not share driver mutations in the majority of tumors.

## Methods

### Mice

BALB/cAnNCrl wild-type and BALB/c CAnN.Cg-*Foxn1^nu^*/Crl athymic mice were purchased from Charles River Laboratories or bred for *in vivo* experiments. Mice were maintained in microisolator cages in the vivarium facility at the University of Arizona College of Medicine – Phoenix in accordance with local and national guidelines. All studies were approved by the University of Arizona’s Institutional Animal Care and Use Committee.

### Cell lines

Solar UV light-induced tumors were generated in BALB/c mice as described,^19^ sterilely harvested, and adapted to cell culture. The central portion of each tumor was used for histological diagnosis and the remaining portions of the tumor were used to create cell lines. Tumor was digested with 2 mg/mL collagenase IV (Worthington Biochemical Corporation Cat. No. LS004188) by shaking for 20 minutes at 37°C, then ground through a 40 μm strainer. The remaining tissue on the strainer, as well as the single cell flowthrough, were separately adapted to cell culture. All cSCC cell lines were cultured in DMEM medium supplemented with 10% fetal bovine serum, 25 mM HEPES, 1% non-essential amino acids, and 1% penicillin-streptomycin. To create clonal cell lines, the cSCC cell lines derived from primary tumors were cloned via limiting dilution. To generate cSCC1.1.1, a wild-type BALB/c mouse was injected with 5×10^6^ cSCC1.1 cells; the tumor was harvested, and cells were adapted to culture and cloned via limiting dilution.

T2-D^d^ and T2-L^d^ cell lines were generously provided by Dr. Peter Cresswell; T2-K^d^ cell lines were generously provided by Dr. Joyce Solheim.^50^ T2-D^d^, T2-K^d^, and T2-L^d^ cell lines were cultured in RPMI medium supplemented with 10% heat-inactivated fetal bovine serum, 15 mM HEPES, and 1% penicillin-streptomycin. T2 cell lines were grown in the presences of 800 μg/mL G418 to select for cells that express the corresponding murine MHC class I allele. All cells tested negative at the start and conclusion of experiments for mycoplasma contamination by PCR.

### Identification of shared mutations in human cSCC

Somatic variants from 88 cSCC tumors across ten datasets were obtained from Chang et al.^18^ Somatic variants from an additional 61 intermediate to high-risk cSCC tumors were obtained from Nassir et al.^20^ These somatic variants were identified using the methods described in the Genome_GPA v5.0.3 algorithm (formerly called TREAT).^51^ To ensure high fidelity calls, the somatic variants from Nassir et al. were further filtered to include only those identified by a consensus of GATK Mutect2 and Strelka2 as described.^16^ Shared mutations were defined as any mutations resulting in the same amino acid change regardless of the specific nucleotide change to account for codon degeneracy. When analyzed separately, the data from Chang et al. had 548 mutations shared between any two tumors, 23 shared between any three tumors, and four shared between any four tumors. A combination of the 27 mutations shared between three and four tumors would cover 53.4% of the total tumors. However, the same 27 mutations only provided 20.45% coverage in the Nassir et al. dataset. When the Nassir et al. dataset was analyzed in isolation, there were 1,015 mutations shared between any two tumors, 40 shared between any three tumors, 12 between any four tumors, and two shared between any five tumors. The 54 mutations shared between three to five tumors would cover 81.97% of the patients, however, the same 54 mutations would only cover 21.31% of the Chang et al. tumors. Therefore, we combined the datasets and restricted to mutations shared in four or more tumors and in at least one tumor from each dataset. Putative driver mutations were annotated using cBioPortal.^52^ The sex, age, immune status, RDEB status, and mutational burden of patients were all compared between patients with and without shared mutations. Multivariate logistic regression modeling was performed to assess the relative contributions of age, RDEB status, and mutational burden on the likelihood of containing a shared mutation. Sex and immune status were not included in the logistic regression model since they were not associated with the likelihood of containing a shared mutation.

### Binding predictions for shared mutations in human cSCC

HLA types were identified for each tumor from the Nassir et al. dataset using Polysolver.^53^ Binding was predicted with NetMHCpan4.0 for patient-specific HLA alleles for the Nassir et al. dataset. Binding mutations were defined as those with at least one 9mer with a predicted binding affinity < 500 nM. For patients for whom we could not determine the HLA types, binding was calculated for the two most common HLA-A, HLA-B, and HLA-C alleles for European individuals, identified from the BeTheMatch registry (https://bioinformatics.bethematchclinical.org/hla-resources/haplotype-frequencies/high-resolution-hla-alleles-and-haplotypes-in-the-us-population/, access date: 8/23/2023).^54^

### RNA/DNA extraction and sequencing

DNA and RNA were extracted from the cSCC cell lines using the DNeasy Blood and Tissue Kit (QIAGEN Cat. No. 69504) and RNeasy Mini kit (QIAGEN Cat. No. 74104), respectively. The manufacturer’s instructions were used to extract DNA and RNA using RNase A (QIAGEN Cat. No. 19101) and DNase I (QIAGEN Cat. No. 79254) treatments, respectively. Whole exome sequencing (WES) at 80x coverage and RNA sequencing (RNAseq) with 20 million read pairs were performed at the Yale Center for Genome Analysis. WES and RNAseq FASTQ files were visualized for quality using FastQC (https://www.bioinformatics.babraham.ac.uk/projects/fastqc/) and trimming was performed with bbduk (<u>sourceforge.net/projects/bbmap/</u>). Sanger sequencing was performed on genomic DNA extracted from cSCC cell lines. All primers were designed using NCBI’s PrimerBLAST.^55^ Primer pairs were designed flanking each target mutation in the cSCC1.1.1 genome by 80-250 base pairs with no predicted off-target loci. If a primer pair specific to the mutation locus could not be designed, then a primer pair was chosen for which the off-target amplicons were of a significantly different (>100 base pairs) size compared to the on-target amplicon. Each mutation locus was amplified in separate PCR reactions using GoTaq Green Master Mix (Promega Cat. No. M7123). PCR reactions were held at 95 °C for two minutes followed by 35 cycles of 45 seconds at 95°C for denaturation, 45 seconds at 58°C for annealing, 30 seconds at 72°C for extension, and five minutes at 72°C for the final extension. Amplified target DNA was purified using QIAquick Gel Extraction Kit (QIAGEN Cat. No. 28704) and sent to the Arizona State University’s Genomics Core for Sanger sequencing.

### Comparison of mutational profile in human and murine cSCC

To ensure that any differences in the human and murine cSCC profile were not attributable to differences in the pipeline, only human data analyzed with a comparable pipeline was used for comparison to the murine cell line. The human data included was that from Nassir et al., Ji et al. (SRP265179),^56^ and Chitsazzadeh et al. (obtained directly from researchers).^57^ The Ji and Chitzazzadeh datasets represent a subset of the data analyzed by Chang et al. that was reanalyzed to ensure accurate comparison to the murine cSCC cell lines. The datasets from Ji and Chitsazzadeh underwent quality control with FastQC (https://www.bioinformatics.babraham.ac.uk/projects/fastqc/), trimming was performed with bbduk (sourceforge.net/projects/bbmap/), and quality was reassessed post-trimming. Read mapping to the 1000 genomes hg38 reference genome^58^ was performed with BWA-mem and read group labels were added using BWA-mem.^59^ SAM files were converted to BAM and coordinate sorted.^60^ Single nucleotide variants (SNVs) and small insertions/deletions (indels) were identified using two programs, along with their recommended filters: GATK Mutect2 version 4.1.7.0^61^ and Strelka version 2.9.2.^62^ To ensure high fidelity calls, these variants were further filtered to include only those identified by a consensus of GATK Mutect2 and Strelka2. Missense mutations were annotated using the Variant Effects Predictor from Ensembl,^63^ and peptides were generated using PVACseq.^64^ The same pipeline was employed to analyze the data from the murine model with the reads aligned to the GRCm38 reference genome from Ensembl.^65^ A matched normal sample from non-solar UV exposed, normal appearing BALB/c ear was used as the reference for each murine cell line.

### Comparison of the driver mutations in human and murine cSCC

A list of known driver genes in cSCC was obtained from Chang et al.^18^ The percentage of human tumors containing a missense or nonsense mutation in a driver mutation was calculated across all data from Nassir et al. and Chang et al. The murine tumors were also queried for missense or nonsense mutations in this set of driver genes.

### Structural modeling

The crystal structure of P53 bound to DNA^24^ was obtained from the Protein Data Bank (PDB code 1TUP)^66^ and the structure of P53 was visualized with Mol* Viewer.^67^ A set of tumor-rejecting neoantigens were identified from existing mouse models. Tumor-rejecting neoantigens from the literature were only included in the analysis if the precise epitope had been individually tested for the ability to elicit tumor-destruction. Structural models of peptide/MHC complexes for all non-immunogenic neoantigens from this study and tumor-rejecting neoantigens from this study and prior studies^9, 11, 12, 17, 27^ were determined using the TFold implementation of Alphafold2 using the previously reported procedure.^28, 68^ Output coordinates were scored as an average of the predicted local-distance difference test (pLDDT) across peptide residues and subtracted from 100. The lowest scores for each output were used to identify models most suitable for subsequent analysis. Overall, confidence scores ranged from 4.01 to 20.66 and 3.67 to 36.54 for negative and positive classes, respectively. We observed that for a subset of the models, the C-terminal residue of the peptide (pΩ) did not properly occupy the pocket of the MHC molecule. Therefore, all models were visually inspected, and those models with pΩ clearly extruding from the groove were excluded from analysis. Solvent accessible surface areas were calculated using the Mol* Viewer using a probe radius of 1.4 Å.^67^ Neoantigens were split into groups based on their DAI, which was calculated as the ratio of the predicted wild-type and mutant peptide dissociation constants. The difference between the SASA of the mutant and corresponding wild-type amino acids was compared between tumor-rejecting neoantigens and non-immunogenic neoantigens in both categories using a Wilcoxon rank sum test. The optimal DAI cut point was selected by comparing a range of cut points between a DAI of 0.6 and 2.5 and selecting the cut point that minimized the p-value for the comparison of tumor-rejecting and non-immunogenic neoantigens in the low DAI category (Extended Data Figure 6).

### Flow cytometry

The following antibodies were used: unconjugated anti-CD16/CD32 (1 μg per 1×10^6^ cells; clone 93; BioLegend Cat. No. 101320), H2-D^d^-FITC (1:200; clone 34-2-12; BioLegend Cat. No. 110606), H2-K^d^-APC (1:160; clone SF1-1.1.1; Invitrogen Cat. No. 17595782), H2-L^d^-PE (1:40; clone 30-5-7S; Invitrogen Cat. No. MA518007), CD45-PerCP-Cy5.5 (1:250; clone 30-F11; Invitrogen Cat. No. 45045182), I-A/E-BV421 (1:80; clone M5/114.15.2; BD Biosciences Cat. No. 562564), CD3-PE-Cy7 (1:80; clone 145-2C11; Invitrogen Cat. No. 25003182), CD8-PE (1:320; clone 53-6.7; BioLegend Cat. No. 100708), CD8-FITC (1:50; clone 53-6.7; BioLegend Cat. No. 100705), CD4-APC (1:80; clone RM4-4; BioLegend Cat. No. 116013), CD4-BV421 (1:160; clone RM4-4; BioLegend Cat. No. 116023), FoxP3-APC (1:20; clone FJK-16s; Invitrogen Cat. No.17577382).

All staining steps were performed at 4°C in the dark. Cells were stained with Fixable Viability Stain 780 (1:2000; BD Biosciences Cat. No. 565388) for 30 minutes. FcγR III/II was blocked with anti-CD16/CD32 antibodies (BioLegend Cat. No. 101320) for five minutes, and conjugated antibodies were added on top of the blocking antibodies for 30 minutes. For intracellular staining, cells were fixed and permeabilized using the True-Nuclear Transcription Factor Buffer Set (BioLegend Cat. No. 424401) using the manufacturer’s protocol with overnight fixation/permeabilization.

The landscape of tumor-infiltrating T cells, spleen-derived T cells, and peripheral T cells from whole blood, was evaluated via flow cytometry. To create a single cell suspension from tumors, tumors were minced and either used fresh, or thawed from previously frozen tissue in a solution of 90% fetal bovine serum and 10% dimethyl sulfoxide. The minced tumor was digested using 2 mg/mL collagenase IV and 20 IU/mL DNase I (Sigma-Aldrich Cat. No. 4716728001) in gentleMACS C tubes (Miltenyi Biotec Cat. No. 130093237) using the gentleMACS Octo Dissociator with Heaters (Miltenyi Biotec Cat. No. 130096427) and the pre-programmed protocol 37C_m_TDK_1. The digestion of tumors was quenched with complete RPMI medium supplemented with fetal bovine serum, 1% non-essential amino acids, and 1% penicillin-streptomycin. Tumors were then ground through a 100 μm strainer twice. To create a single cell suspension from spleens, spleens were mechanically dissociated through a 100 μm strainer. Submandibular blood was collected from live mice and immediately placed in 4% citric acid on ice to avoid coagulation. Blood, and the single cell suspensions from tumors and spleens, were incubated with ACK buffer (0.15 M NH_4_Cl, 1 M KHCO_3_, 0.1 mM Na_2_EDTA) for ten and five minutes, respectively, to lyse red blood cells. After treatment with ACK buffer, all samples were filtered through a 40 μm strainer and stained as indicated above.

The expression of BALB/c-specific MHC class I alleles was evaluated *in vitro* for cell lines and *ex vivo* in mouse tumors. T2 cells transfected with BALB/c-specific MHC class I alleles, cSCC cell lines, and cSCC cell line-derived mouse tumors that did not respond to treatment with vaccination, were stained with H2-D^d^-FITC, H2-K^d^-APC, and H2-L^d^-PE antibodies using the methods stated above. To enrich the population of tumor cells from the non-adherent leukocytes, digested *ex vivo* tumors were grown in TC-treated plates overnight prior to staining, and trypsinized the day of staining to create a single cell suspension of the adherent cells; additionally, cells were stained with CD45-PerCP-Cy5.5, and tumor cells were gated on CD45-negative.

Flow cytometry was performed on the LSRII (BD Biosciences) or NovoCyte Quanteon (Agilent) and analysis was performed on FlowJo v10.7.1 (BD Biosciences) or the NovoExpress software (Agilent). Prior to gating the cell populations of interest, samples were first gated on FSC vs. SSC to exclude debris, FSC-A vs. FSC-H to exclude doublet cells, and Fixable Viability Stain 780 to exclude dead cells. Precision Count Beads (BioLegend Cat. No. 424902) were used to establish the absolute count of cells, or these data were derived from the NovoExpress software. Fluorescence minus one controls were used when needed and single stained controls of cells or UltraComp eBeads (Invitrogen Cat. No. 01222242) were used for compensation.

### Tumor transplantation

To establish tumors, 100 μL of cells were injected intradermally into the right flank of six- to ten-week-old male wild-type or athymic BALB/c mice. Since cSCC1.1, cSCC1.2, cSCC1.3, cSCC1.1.1, and cSCC3.1 were derived from tumors from male mice, only male mice were used to prevent the recognition of Y-encoded antigens by the immune system of female mice.^69^ Since cSCC2.1 and cSCC2.2 derived from a tumor in a female mouse, both male and female mice were used for *in vivo* experiments. Tumors were measured thrice weekly or daily and the tumor volume was calculated using the formula volume = (L*W*H*π)/6.^70^ For tumors measured daily, mice were euthanized after the first tumor volume measurement ≥ 2000 mm^3^. For tumors measured thrice weekly, mice were euthanized after the first tumor volume measurement ≥ 1500 mm^3^.

### Antibody depletions

For depletion studies, mice were weighed prior to each treatment with depleting antibodies. Mice were intraperitoneally injected with 10 mg/kg of anti-CD8α (clone 2.43; Bio X Cell Cat. No. BE0061) and/or anti-CD4 (clone GK1.5; Bio X Cell Cat. No. BE0003-1), to deplete CD8 and CD4 T cells, respectively. Control mice were intraperitoneally injected with 10 mg/kg of rat IgG2b isotype control (clone LTF-2; Bio X Cell Cat. No. BE0090). Unless stated otherwise, mice were treated with depleting antibodies on days −1 and +3 relative to tumor challenge, and weekly thereafter. One day prior to the initiation of treatment with depleting antibodies, blood was taken from each mouse and stained for flow cytometry to establish a baseline level of CD8 and CD4 T cells. Approximately two weeks after the baseline levels of CD8 and CD4 T cells were established, blood was taken from each mouse to confirm depletion of the desired T cell population(s). Once mice had tumors that reached the tumor volume endpoint, tumors and/or spleens were harvested and used to confirm depletion of the desired T cell population(s), following the methods detailed in the “Flow cytometry” section.

### *In vitro* doubling time

cSCC1.1 and cSCC1.1.1 cells were each grown in four separate, clear, flat-bottom 96 well plates. Plates were harvested every 24 hours for 96 hours total, with one plate per cell line read at each timepoint. CellTiter-Glo 2.0 Cell Viability Assay (Promega Cat. No. G9242) was used according to the manufacturer’s instructions as a measure of proliferation. Luminescence was read on a SafireII Microplate Reader (Tecan), once supernatant was transferred to a white-walled 96 well plate. Each well was done in triplicate and luminescence was normalized to media alone controls.

### Peptides

All 9mer and 21mer neoantigen peptides used for *in vitro* and *in vivo* experiments were purchased from GenScript (Piscataway, NJ). For 21mer neoantigen peptides, the mutation was centered within the peptide. The neoantigen peptides were high-performance liquid chromatography purified to at least 95% purity, and the sequences were confirmed with mass spectrometry. Trifluoroacetic acid was removed, so that each neoantigen peptide contained less than 1% trifluoroacetic acid. Mycobacterium tuberculosis peptide (GGPHAVYLL), which binds H2-D^d^; influenza hemagglutinin peptide (IYSTVASSL), which binds H2-K^d^; and β-galactosidase peptide (TPHPARIGL), which binds H2-L^d^, were used as control peptides in *in vitro* and *in vivo* experiments.

### Vaccinations

Cultured cells resuspended in PBS were irradiated with 10-80 Gy using an X-RAD 320 irradiator (Precision X-Ray Inc). After irradiation, cells were intradermally injected into the left flank of mice. Mice were vaccinated with irradiated cSCC cells 14 and 7 days, or 7 days, prior to tumor challenge. For prophylactic vaccination with peptides, mice were intradermally injected with 50 μg of neoantigen or control peptide and/or 100 μg high molecular weight VacciGrade polyinosinic-polycytidylic acid (poly(I:C) (Invivogen Cat. No. vac-pic)). All mice vaccinated with peptides received two doses 14 and 7 days prior to tumor challenge.

### Neoantigen prioritization in murine cSCC cell lines

To select a list of high confidence, prioritized immunogenic neoantigens in the murine cSCC cell lines, somatic variants were first confirmed by presence in at least one RNAseq replicate using GATK ASEReadCounter.^61^ Neoantigens were then prioritized based on predicted MHC binding and variant allele-specific mRNA expression. MHC binding was predicted using the NetMHCpan4.1 suite.^26^ Neoantigens were prioritized with a combination of binding affinity percent rank, which predicts the interaction between HLA alleles and identified neoantigens using IC50 data, and the eluted ligand percent rank, which predicts the likelihood of neoantigens from being eluted from HLA alleles using mass spectrometry. Neoantigens were filtered for a binding affinity percent rank and eluted ligand percent rank under 0.5% for MHC class I alleles. For the RNA expression, RNAseq data underwent transcriptome assembly and read count quantifications with Salmon version 0.11.3,^71^ using the GRCm38 reference transcriptome. mRNA expression was quantified in units of transcripts per million and averaged across all replicates for each sample. Variant allele-specific expression was quantified by multiplying transcript-level expression by the ratio of variant reads to total reads as quantified by GATK ASEReadCounter.

### ELISPOT

Tumors and tumor-draining lymph nodes (TDLN) were harvested from mice ten days after tumor challenge. The axillary, brachial, cervical, inguinal, and mandibular lymph nodes (LN) and spleens were harvested from naïve age-matched mice. Tumors were minced and digested using collagenase IV as described in “Flow cytometry” section to obtain a single cell suspension, though DNase I was not used for ELISPOT tumor digestion. For TDLN, naïve LN, and spleens, the tissues were mechanically dissociated through a 100 μm strainer. Splenocytes were incubated with ACK buffer for five minutes to lyse red blood cells. After five minutes, the lysis reaction was quenched with T cell media (RPMI medium supplemented with 10% heat-inactivated fetal bovine serum, 1% penicillin-streptomycin, 1% non-essential amino acids, 1% sodium pyruvate, 55 µM 2-mercaptoethanol, and 100 IU/mL IL-2 (PeproTech Cat. No. 21212; Cranbury, NJ)). TDLN, naïve LN, and splenocytes were filtered again through a 40 µm strainer prior to use with isolating beads. Live cells from tumors, TDLN, and naïve LNs, were first isolated using the negative selection Annexin V Dead Cell Removal Kit (STEMCELL Cat. No. 17899). Then, CD4 T cells were isolated using the Mouse CD4 Positive Selection Kit II (STEMCELL Cat. No. 18952), and the flowthrough was used to isolate CD8 T cells using the Mouse CD8α Positive Selection Kit II (STEMCELL Cat. No. 18953). All EasySep isolation kits were used according to the manufacturer’s instructions. For T cells isolated from tumors, 1.25×10^4^ CD8 or CD4 T cells, 6.25×10^4^ naïve splenocytes (APCs), and 2 µg/mL of peptide, were co-cultured in T cell media. For T cells isolated from TDLN and naïve LNs,1.25×10^5^ CD8 or CD4 T cells, 6.25×10^5^ naïve APCs, and 2 µg/mL of peptide were co-cultured in T cell media. T cells and splenocytes treated with 50 ng/mL of PMA (LC Laboratories Cat. No. P1680) and 1 μg/mL of ionomycin (Pepro Tech Cat. No. 5608212) served as a positive control. Negative controls included: T cells and APCs without peptide; T cells alone; and T cells and APCs with irrelevant control peptides that bind each of the three BALB/c-specific MHC class I alleles. All cells were co-cultured for 48 hours in mouse IFN-γ pre-treated ELISPOT plates, and ELISPOT plates were developed following the manufacturer’s protocol for the ELISpot Plus: Mouse IFN-γ (ALP) kit (Mabtech Cat. No. 3321-4APW). After the plates were developed, they were left to dry overnight, and spots were quantified using an AID Classic ELISPOT reader (Autoimmun Diagnostika).

### RT-qPCR Methods

One or two sections of cSCC1.1.1. tumors from poly(I:C) or mKars vaccinated mice were flash-frozen. For RNA extraction, tumor sections were first homogenized using tissue grinding tubes (Bertin Corporation Cat. No. P000922-LYSK0-A), and three iterations of grinding at 6500 rpm for 20 seconds, followed by 30 seconds on ice, on the Precellys 24 Tissue Homogenizer. RNA was extracted from the homogenized tissue, as well as cultured cSCC1.1.1 cells, as detailed in “RNA/DNA extraction and sequencing.” *Kars* RNA expression was measured in each tumor using the iTaq Universal SYBR Green One-Step Kit (Bio-Rad Cat. No. 1725150) and the Quantstudio 6 Flex System (Applied Biosystems). *Kars* RNA was reverse-transcribed then amplified using primers specific to an 85 bp region overlapping the junction of exon six and seven in the *Kars* gene. Primers were designed using PrimerBLAST.^72^ *Gapdh* RNA was used as an endogenous control and amplified using primers as described.^73^ Reactions were held at 50°C for ten minutes, then at 95°C for one minute, followed by 40 cycles of 15 seconds at 95°C for denaturation, and 30 seconds at 60°C for annealing and extension. Cycling was then followed by a melt curve analysis to confirm that amplification was target-specific. All reactions were performed in triplicate.

### MHC class I stabilization and relative binding affinity using T2 cells

The stabilization of BALB/c-specific MHC class I alleles due to peptide binding was determined using T2 cells, as previously described.^50^ In brief, T2 cells were harvested, plated at 2×10^5^ cells per well, and incubated for 24 hours at 26°C. After 24 hours, each well was treated with 10 µg/mL brefeldin A (Medchemexpress LLC Cat. No. HY16592) and 100 µg/mL of peptide. Plates were then incubated at 37°C for one, two, or four hour(s). Cells at the zero hour timepoint were not treated with brefeldin A or incubated with peptide. To determine the relative peptide:MHC binding affinities, similar to above, T2 cells were incubated for 24 hours at 26°C, and then treated with brefeldin A and a peptide titration for four hours at 37°C. In both cases, cells were kept on ice following the completion of the 37°C incubation. Cells were stained for cell surface MHC class I expression following the methods detailed in “Flow cytometry.”

### Statistical analysis

Statistical analysis was performed with *R* software version 4.1.0 (R Core Team) or Prism 9 software (GraphPad).

## Data availability

RNAseq and WES data for all murine tumors and cell lines will be uploaded to the Sequencing Reads Archive (SRA) upon manuscript acceptance.

## Code availability

Code for the analysis of the human samples is available at https://github.com/HastingsLab/Human_SCC_data and code for the analysis of the murine samples is available at https://github.com/HastingsLab/SSL_SSC_Neoantigens.

## Acknowledgments

This work was supported in part by National Institutes of Health grants T32CA009213 (A.C.A.), F30CA257622 (A.C.A.), F30CA281056 (E.S.B.), P30CA023074 (D.J.R.), R35GM118166 (B.M.B.), and P01CA229112 (K.T.H. and D.J.R.); grant from the American Skin Association (A.C.A.); University of Arizona grants Cancer Center Training Oncology Physician Scientists Award (E.S.B), Springboard Initiative (K.T.H.), and Grand Challenges in Healthy Aging (K.T.H.); Dermatology Foundation Career Development Award (A.R.M.), Mayo Clinic Investment for Extramural Grants Award (A.R.M.), Mayo Clinical Cancer Center Desert Mountain Care program (A.R.M.), and Arnold and Kit Palmer Career Development Award in Cancer Research (A.R.M.). This work benefited from support for a related project by the Merit Review Award I01-BX005336 from the United States Department of Veterans Affairs (VA), Biomedical Laboratory Research and Development Service (K.T.H.). The contents do not represent the views of the VA or the United States Government. We thank Karen S. Anderson and Padhmavathy Yuvaraj for training and use of the ELISPOT reader. We thank dermatopathologist David J. Glembocki for histologically diagnosing the tumors used to generate cSCC cell lines. The mouse schematic in Figure 3 was created using BioRender.

## Author contributions

Cell lines were generated by A.C.A. and A.M.M. under the guidance of K.T.H. A.C.A., A.M.M., and K.T.H. designed experiments. A.C.A., A.M.M., and L.M.H. completed experiments. E.S.B. completed all human and mouse bioinformatic analyses and prioritized neoantigens with guidance from K.H.B., M.A.W., and K.T.H. T.M., X.L., A.H., and A.R.M. provided the human cSCC dataset from Mayo Clinic. C.A.B. and B.M.B. completed structural modeling. A.C.A., E.S.B., C.A.B., B.M.B., and K.T.H. evaluated features of tumor-rejecting neoantigens. A.C.A. and D.J.R. analyzed data and performed statistical analyses. A.C.A. prepared the figures. A.C.A., A.M.M., E.S.B., B.M.B., and K.T.H. wrote the manuscript with input from each author. A.C.A. and K.T.H. obtained funding for this project. K.T.H. oversaw the project.

---

**Extended Data Figure 1:**
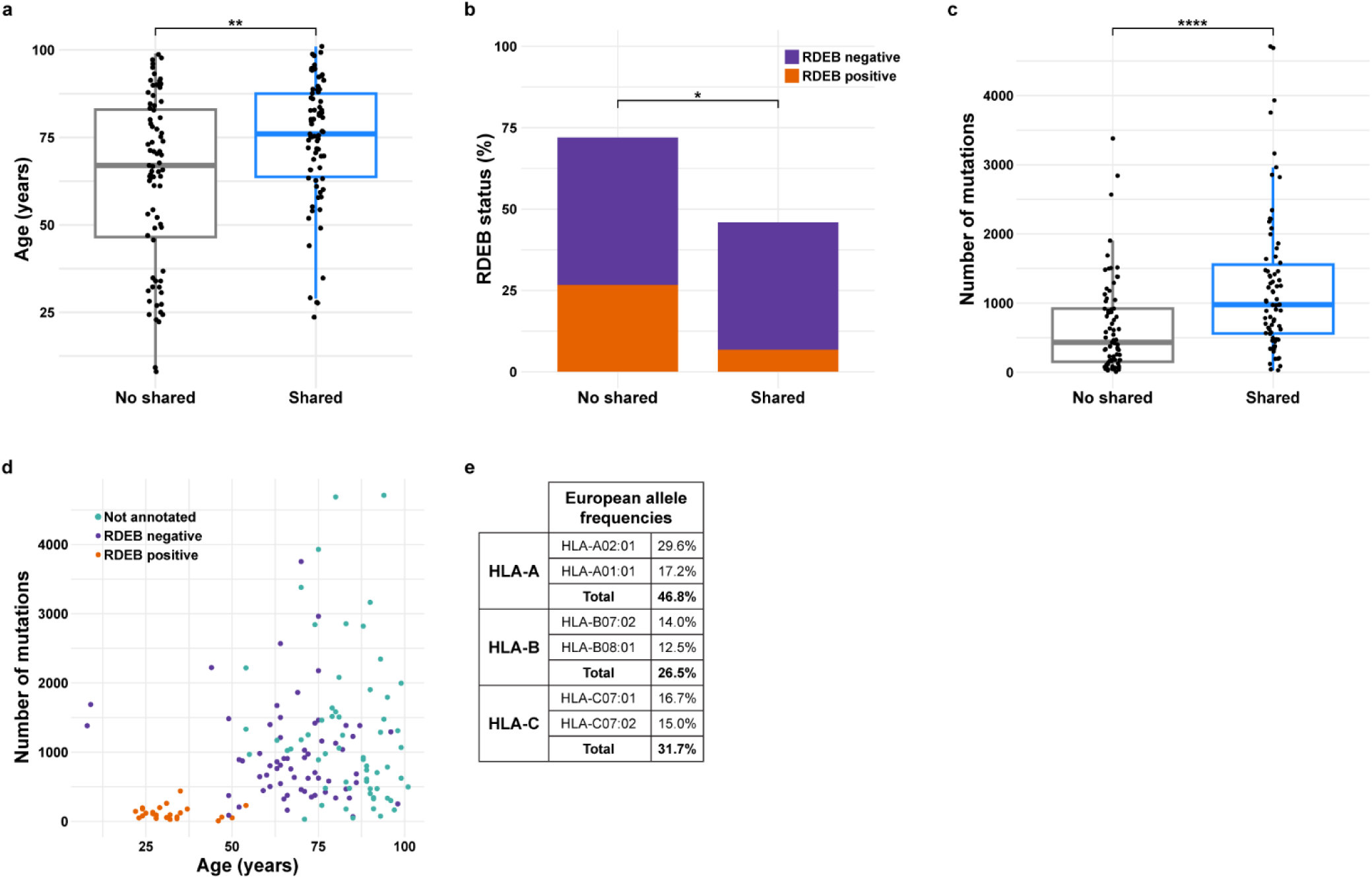
Demographic characteristics of patients with shared mutations. **a,** Box plot displaying the age distribution of patients with cSCC tumors with and without shared mutations. **b,** Stacked bar graph displaying the percentage of patients with and without shared mutations that have recessive dystrophic epidermolysis bullosa (RDEB). Tumors for which the RDEB status was not annotated are not displayed. **c,** Box plot displaying the distribution of the total number of missense mutations in tumors with and without shared mutations. **d,** Plot displaying the number of mutations versus patient age, colored by RDEB status. **e,** Frequency of the most common HLA-A, HLA-B, and HLA-C alleles in individuals of European descent. For box plots, the bold line indicates the median, and the upper and lower limits of the boxes indicate the 75th and 25th percentiles, respectively. The lower and upper whiskers indicate the minimum and maximum. Dots outside of the box and whiskers indicate outliers. Statistical significance was assessed by Wilcoxon rank sum test (**a, c**) or chi-squared test (**b**). *p<0.05; **p < 0.01; ****p < 0.0001

**Extended Data Figure 2:**
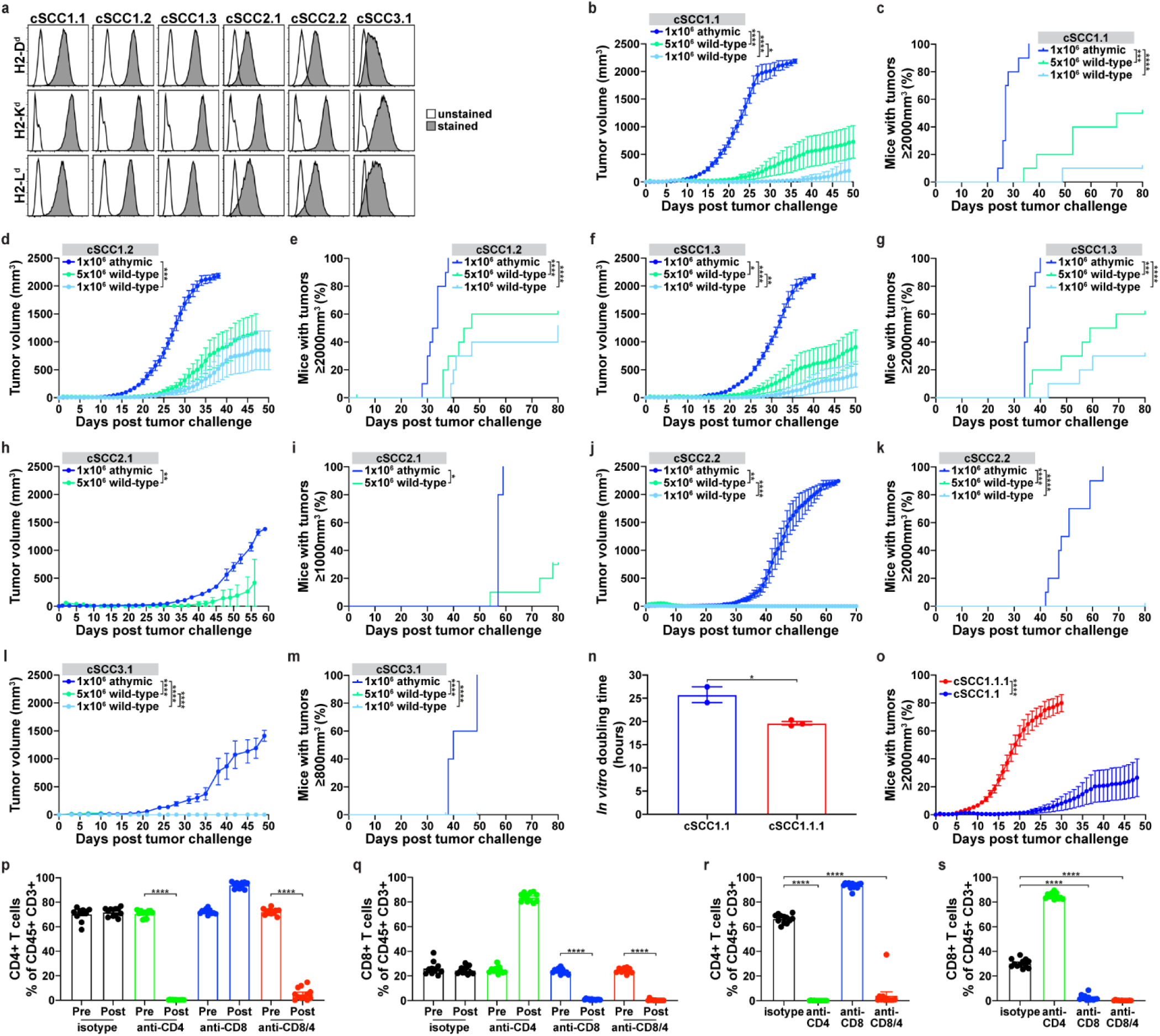
Panel of immunogenic cSCC cell lines constrained by T cells. **a,** *In vitro* expression of BALB/c-specific MHC class I alleles in cSCC cell lines using flow cytometry. Tumor volume and survival curves for mice intradermally injected with **b-c,** cSCC1.1; **d-e,** cSCC1.2; **f-g,** cSCC1.3; **h-i,** cSCC2.1; **j-k,** cSCC2.2; and **l-m,** cSCC3.1. Data pooled from two to three independent experiments (n = 5 - 10 mice per group). **n,** *In vitro* doubling time for cSCC1.1 and cSCC1.1.1 was compared using an unpaired Student’s t test. **o,** Tumor volume curves for wild-type mice intradermally injected with either 2.5×10^6^ cSCC1.1 (n = 10) or cSCC1.1.1 (n = 20) cells. Data pooled from multiple independent experiments. **p-q,** Confirmation of CD4 and/or CD8 T cell depletion in blood for each mouse in the experiment described in Fig. 3f-h. The percentages of CD8 and CD4 T cells before and after the initiation of depleting antibodies were compared in blood using a paired Student’s t test. **r-s,** Once mice reached the tumor volume endpoint or the experiment ended, spleens were harvested from mice described in Fig. 3f-h, and were used to confirm depletion of targeted T cell populations. The percentages of CD4 or CD8 T cells in spleens for mice treated with depleting antibodies were compared to mice treated with isotype control using an unpaired Student’s t test. Tumor volume (mean ± SEM) over time displayed using last observation carried forward. Tumor growth was compared using a linear mixed effects model and time to reach tumor volume endpoint was compared using a Cox proportional hazards model. *p<0.05; **p < 0.01; ***p < 0.001; ****p < 0.0001

**Extended Data Figure 3:**
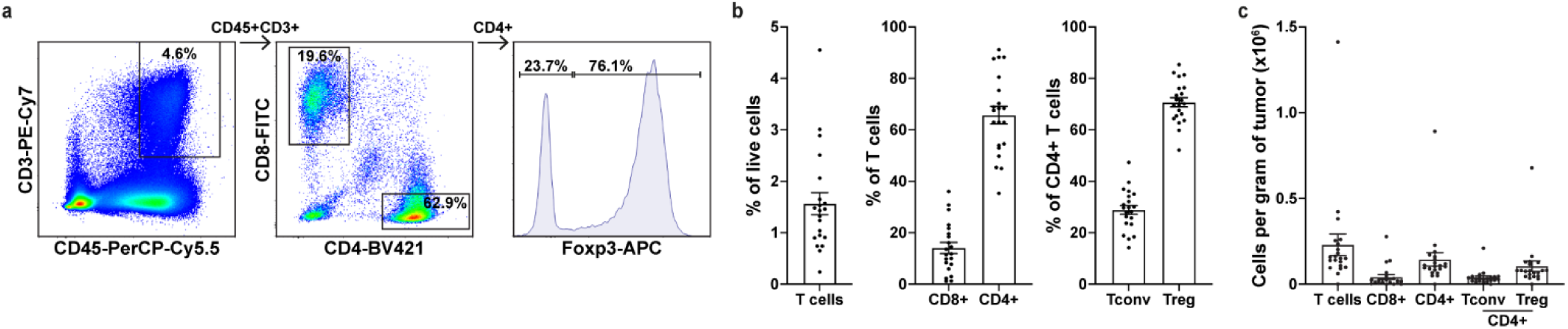
The majority of T cells in cSCC1.1.1 tumors are CD4 T regulatory cells. **a,** Gating strategy for the identification of T cell populations within cSCC1.1.1 tumors that reached the tumor volume endpoint. **b,** Box plots showing the percentage of intratumoral T cells out of live cells (left panel), the percentage of CD8+ and CD4+ T cells out of CD45+ CD3+ T cells (middle panel), and the percentage of conventional (Tconv) and regulatory (Treg) cells out of CD4+ T cells (right panel) in cSCC tumors. **c,** Box plot showing the absolute count of T cells, CD8+ T cells, CD4+ T cells, CD4+ Tconv cells, and CD4+ Treg cells per gram of tumor. Percentage and absolute count of T cells (mean ± SEM) from n = 21 cSCC1.1.1 tumors.

**Extended Data Figure 4:**
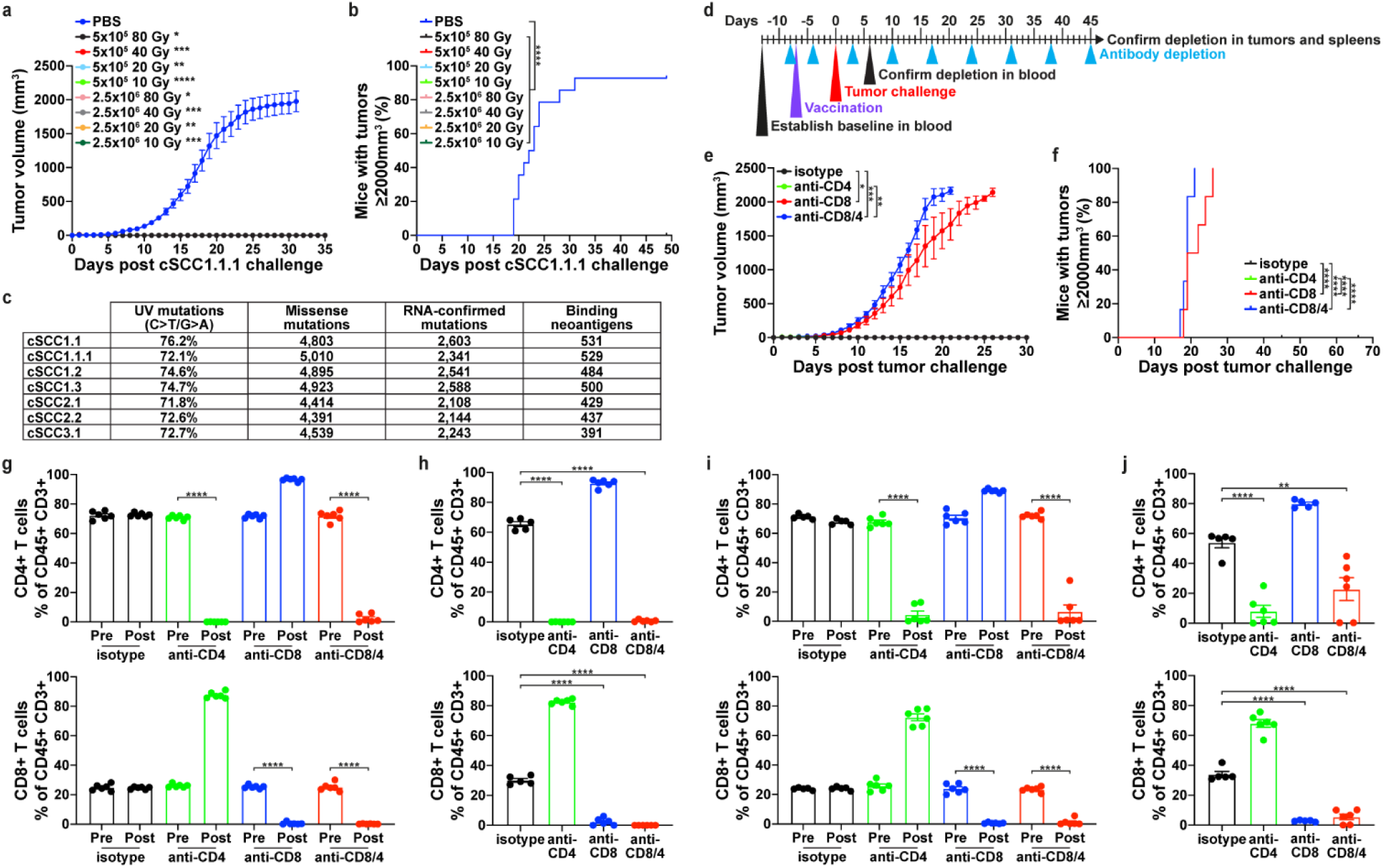
CD8 T cells are responsible for response to vaccination with irradiated tumor cells. **a,** Wild-type mice were intradermally vaccinated with PBS, 5×10^5^ cSCC1.1.1 cells, or 2.5×10^6^ cSCC1.1.1 cells irradiated with 10-80 Gy. One week after vaccination, mice were intradermally injected with 2.5×10^6^ cSCC1.1.1 cells. Data shown are from two pooled experiments with n = 6 - 14 mice per group. **b,** Survival curves for mice described in (**a**). **c,** UV-signature mutations (C>T/G>A), missense mutations, RNA-confirmed mutations, and binding MHC class I neoantigens in the panel of clonal cSCC cell lines. **d,** Schematic of vaccination and T cell depletion strategy for the priming phase of vaccination. Wild-type mice were intradermally vaccinated with 5×10^5^ cSCC1.1.1 cells irradiated with 10 Gy on day −7 relative to tumor challenge. Mice received intraperitoneal treatment with isotype control, anti-CD4, anti-CD8, or anti-CD8/4 on days −8 and −4 relative to intradermal injection with 2.5×10^6^ cSCC1.1.1 cells, and weekly thereafter. **e,** Tumor volume curves and **f,** survival curves for mice described in (**d**). Data shown are from one experiment with n = 6 mice per group. **g,** For the mice described in (**d**), blood was taken from each mouse before and after treatment with T cell depleting antibodies. T cell populations were evaluated using flow cytometry. **h,** At the end of the experiment described in (**d**) or once tumor-bearing mice reached the tumor volume endpoint, spleens were harvested from each mouse. Depletion of CD8 and/or CD4 T cell populations were confirmed using flow cytometry. **i,j,** Confirmation of CD8 and/or CD4 T cell populations for mice described in Fig. 4i-k in blood throughout the experiment (**i**) and in spleens as mice reached the tumor volume endpoint or the end of the experiment (**j**). All tumors were measured daily, and mice were euthanized when tumors were ≥ 2000 mm^3^. Tumor volume (mean ± SEM) over time displayed using last observation carried forward. Tumor growth was compared using a linear mixed effects model, and time to reach tumor volume endpoint was compared using a Cox proportional hazards model. The percentage of CD8 and CD4 T cells before and after depletion was compared in blood using a paired Student’s t test. The percentage of CD8 and CD4 T cells in spleens was compared using an unpaired Student’s t test. *p<0.05; **p < 0.01; ***p < 0.001; ****p < 0.0001

**Extended Data Figure 5:**
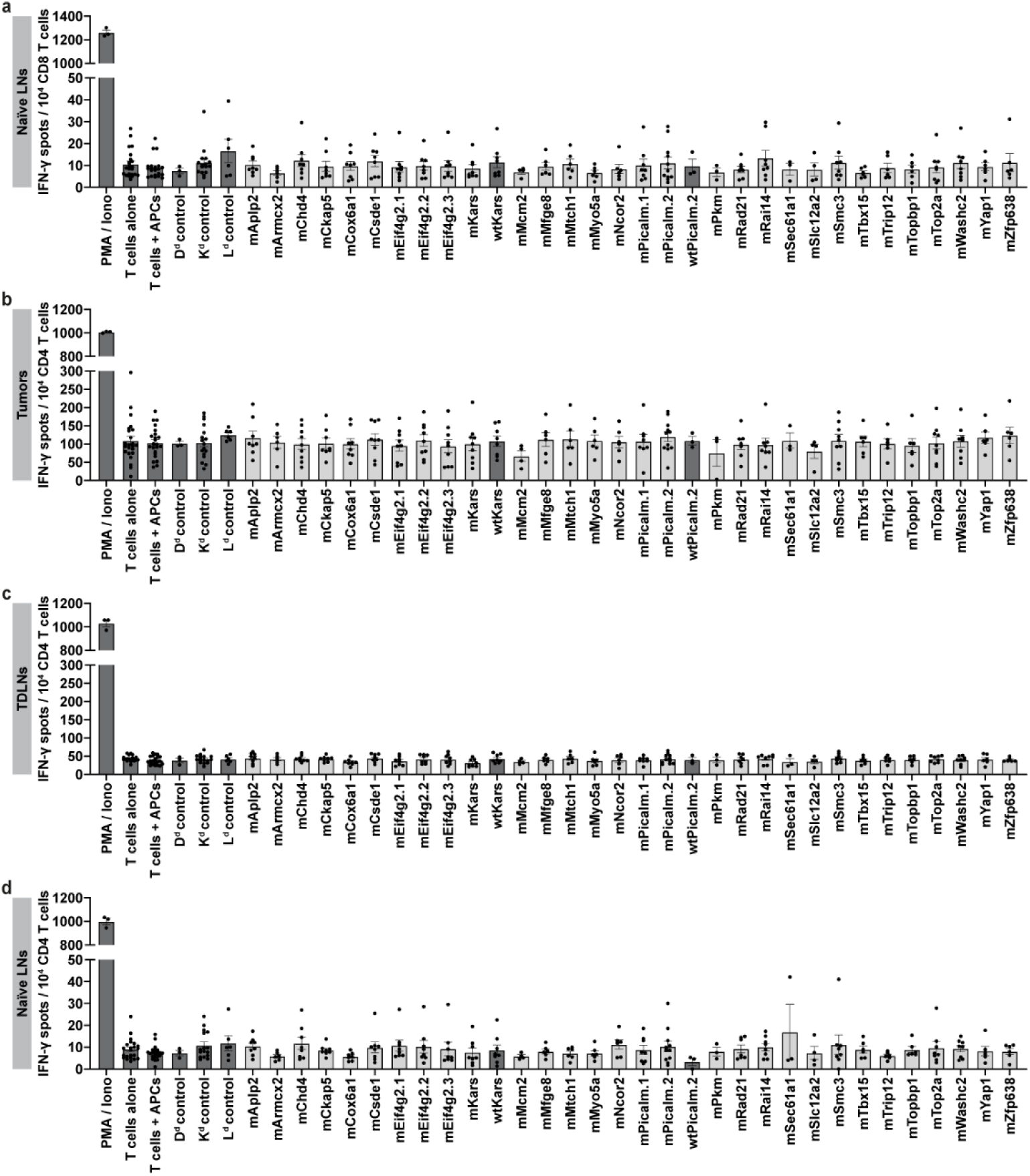
Predicted MHC class I neoantigens do not elicit CD4 T cell responses in tumor challenged mice and do not elicit T cell responses in naïve mice. Ten days after mice were challenged with 2.5×10^6^ cSCC1.1.1 cells, tumors and tumor draining lymph nodes (TDLNs) were harvested. Lymph nodes (LNs) were harvested from naïve mice. T cells isolated from mice were used for IFN-γ enzyme-linked immune absorbent spot (ELISPOT). ELISPOT results for **a,** CD8 T cells isolated from naïve LNs; **b,** CD4 T cells isolated from tumors; **c,** CD4 T cells isolated from TDLNs; and **d,** CD4 T cells isolated from naïve LNs. Isolated T cells were co-cultured with naïve splenocytes (APCs) pulsed with 2 µg/mL of the indicated individual peptides. Data displayed as mean ± SEM (n = 3 - 9 independent experiments). IFN-γ spots for all negative control groups (T cells alone; T cells + APCs; and irrelevant binding peptides: D^d^ control, K^d^ control, and L^d^ control) were pooled and compared to each mutant (m) peptide using a Kruskal-Wallis test. Mutant peptides were compared to wild-type (wt) peptides using a Mann-Whitney test.

**Extended Data Figure 6:**
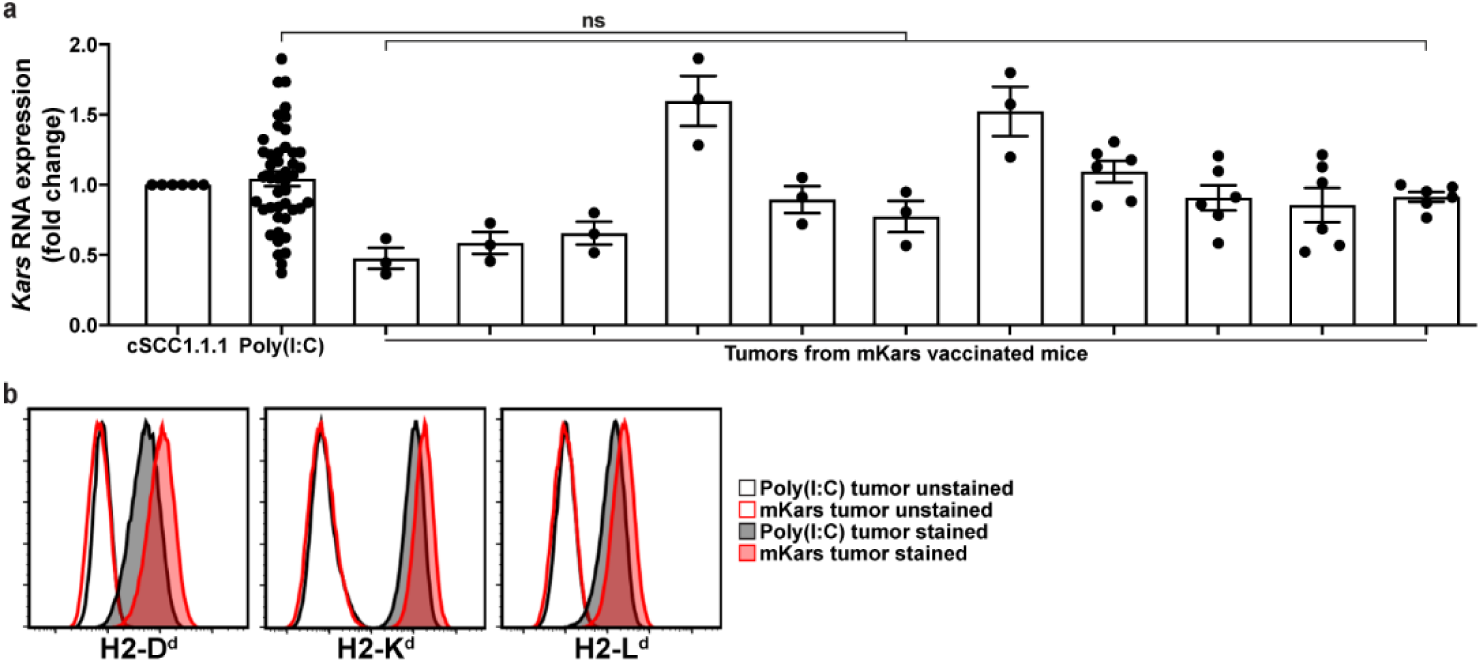
Tumors that escape vaccination with mKars maintain *Kars* RNA expression and tumor cell expression of MHC class I. **a,** Tumors from poly(I:C) vaccinated mice or 9mer mKars vaccinated mice were harvested, and the RNA expression of *Kars* was evaluated by qPCR. *Kars* RNA expression in tumors from poly(I:C) vaccinated mice are pooled together in the display, and expression in tumors from 9mer mKars vaccinated mice are shown individually. *Kars* RNA expression was normalized to *Gapdh* and expressed as fold change relative to the *in vitro* expression in cSCC1.1.1 using the comparative Ct method (mean ± SEM) (n = 11 tumors per group, one or two sections per tumor, each section performed in triplicate). *Kars* RNA expression was compared between each tumor from an mKars vaccinated mouse and the pooled tumors from poly(I:C) treated mice using an unpaired Student’s t test. **b,** Tumors from mice vaccinated with poly(I:C) or 9mer mKars were cultured overnight, and adherent tumor cells were evaluated for expression of MHC class I alleles using flow cytometry. Tumor cells were considered CD45 negative. Plots are representative of all tumors tested (n = 11 mice per group). ns (not significant)

**Extended Data Figure 7:**
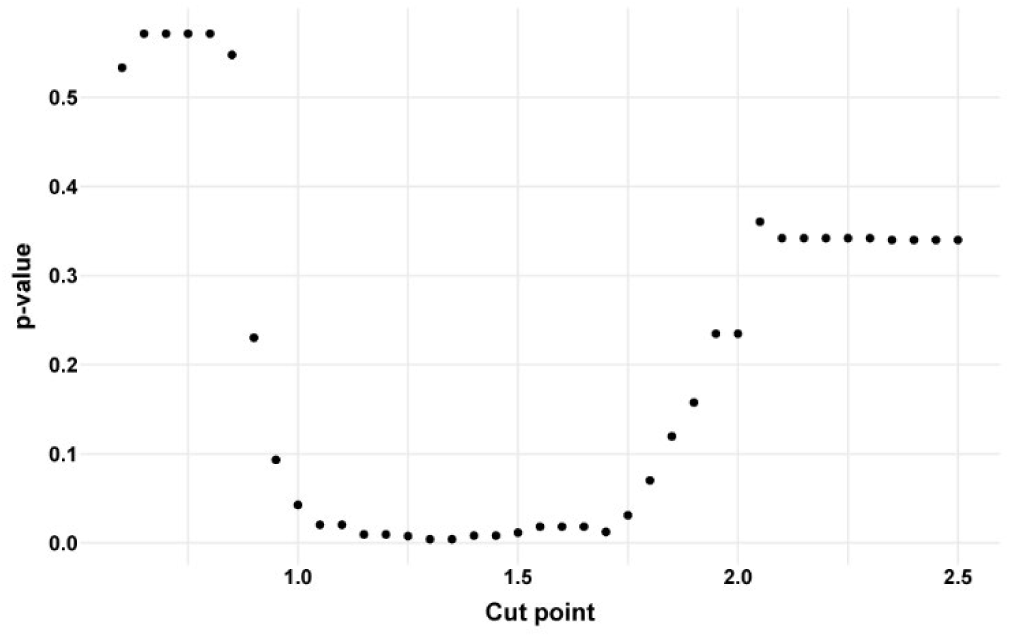
Optimization of differential agretopic index (DAI) cut point. Neoantigens were separated into a DAI high and DAI low class using a range of thresholds from 0.6 to 2.5 with 0.05 increments. At each increment, the difference between the mutant and wild-type SASA was compared between tumor-rejecting neoantigens and non-immunogenic neoantigens in the low DAI class, as described in Fig. 7. The p-value was plotted against the cut point to select an optimal cut point.

## References

1. Schumacher, T.N. & Schreiber, R.D. Neoantigens in cancer immunotherapy. Science 348, 69–74 (2015).

2. Rojas, L.A. et al. Personalized RNA neoantigen vaccines stimulate T cells in pancreatic cancer. Nature 618, 144–150 (2023).

3. Sahin, U. et al. Personalized RNA mutanome vaccines mobilize poly-specific therapeutic immunity against cancer. Nature 547, 222–226 (2017).

4. Ott, P.A. et al. An immunogenic personal neoantigen vaccine for patients with melanoma. Nature 547, 217–221 (2017).

5. Hilf, N. et al. Actively personalized vaccination trial for newly diagnosed glioblastoma. Nature 565, 240–245 (2019).

6. Weber, J.S. et al. Individualised neoantigen therapy mRNA-4157 (V940) plus pembrolizumab versus pembrolizumab monotherapy in resected melanoma (KEYNOTE-942): a randomised, phase 2b study. The Lancet.

7. Katsikis, P.D., Ishii, K.J. & Schliehe, C. Challenges in developing personalized neoantigen cancer vaccines. Nature Reviews Immunology (2023).

8. Carreno, B.M. et al. Cancer immunotherapy. A dendritic cell vaccine increases the breadth and diversity of melanoma neoantigen-specific T cells. Science 348, 803–808 (2015).

9. Gubin, M.M. et al. Checkpoint blockade cancer immunotherapy targets tumour-specific mutant antigens. Nature 515, 577–581 (2014).

10. Castle, J.C. et al. Exploiting the mutanome for tumor vaccination. Cancer Res 72, 1081–1091 (2012).

11. Duan, F. et al. Genomic and bioinformatic profiling of mutational neoepitopes reveals new rules to predict anticancer immunogenicity. Journal of Experimental Medicine 211, 2231–2248 (2014).

12. Dolina, J.S. et al. Linked CD4+/CD8+ T cell neoantigen vaccination overcomes immune checkpoint blockade resistance and enables tumor regression. J Clin Invest 133 (2023).

13. Yadav, M. et al. Predicting immunogenic tumour mutations by combining mass spectrometry and exome sequencing. Nature 515, 572–576 (2014).

14. Alspach, E. et al. MHC-II neoantigens shape tumour immunity and response to immunotherapy. Nature 574, 696–701 (2019).

15. Wells, D.K. et al. Key Parameters of Tumor Epitope Immunogenicity Revealed Through a Consortium Approach Improve Neoantigen Prediction. Cell 183, 818–834.e813 (2020).

16. Borden, E.S. et al. NeoScore Integrates Characteristics of the Neoantigen:MHC Class I Interaction and Expression to Accurately Prioritize Immunogenic Neoantigens. J Immunol (2022).

17. Ebrahimi-Nik, H., et al. Mass spectrometry driven exploration reveals nuances of neoepitope-driven tumor rejection. JCI Insight 5 (2019).

18. Chang, D. & Shain, A.H. The landscape of driver mutations in cutaneous squamous cell carcinoma. npj Genomic Medicine 6, 61 (2021).

19. Adams, A.C. et al. Solar simulated light induces cutaneous squamous cell carcinoma in inbred mice: a clinically relevant model to investigate T cell responses. J Invest Dermatol (2021).

20. Nassir, S. et al. Whole exome and transcriptome sequencing of stage-matched, outcome-differentiated cutaneous squamous cell carcinoma identifies gene expression patterns associated with metastasis and poor outcomes. medRxiv, 2024.2002.2005.24302298 (2024).

21. Lawrence, M.S. et al. Discovery and saturation analysis of cancer genes across 21 tumour types. Nature 505, 495–501 (2014).

22. Hoyos, D. et al. Fundamental immune-oncogenicity trade-offs define driver mutation fitness. Nature 606, 172–179 (2022).

23. Jorgenson, E. et al. Genetic ancestry, skin pigmentation, and the risk of cutaneous squamous cell carcinoma in Hispanic/Latino and non-Hispanic white populations. Communications Biology 3, 765 (2020).

24. Emamzadah, S., Tropia, L. & Halazonetis, T.D. Crystal Structure of a Multidomain Human p53 Tetramer Bound to the Natural CDKN1A (p21) p53-Response Element. Molecular Cancer Research 9, 1493–1499 (2011).

25. Jurtz, V. et al. NetMHCpan-4.0: Improved Peptide-MHC Class I Interaction Predictions Integrating Eluted Ligand and Peptide Binding Affinity Data. J Immunol 199, 3360–3368 (2017).

26. Reynisson, B., Alvarez, B., Paul, S., Peters, B. & Nielsen, M. NetMHCpan-4.1 and NetMHCIIpan-4.0: improved predictions of MHC antigen presentation by concurrent motif deconvolution and integration of MS MHC eluted ligand data. Nucleic Acids Research 48, W449–W454 (2020).

27. Brennick, C.A. et al. An unbiased approach to defining bona fide cancer neoepitopes that elicit immune-mediated cancer rejection. The Journal of Clinical Investigation 131 (2021).

28. Mikhaylov, V. et al. Accurate modeling of peptide-MHC structures with AlphaFold. Structure (2023).

29. Custodio, J.M. et al. Structural and physical features that distinguish tumor-controlling from inactive cancer neoepitopes. Proc Natl Acad Sci U S A 120, e2312057120 (2023).

30. Schmidt, J. et al. Prediction of neo-epitope immunogenicity reveals TCR recognition determinants and provides insight into immunoediting. Cell Rep Med 2, 100194 (2021).

31. Chowell, D. et al. TCR contact residue hydrophobicity is a hallmark of immunogenic CD8+ T cell epitopes. Proc Natl Acad Sci U S A 112, E1754–1762 (2015).

32. Calis, J.J.A. et al. Properties of MHC Class I Presented Peptides That Enhance Immunogenicity. PLOS Computational Biology 9, e1003266 (2013).

33. Zhang, H., Zhong, A. & Chen, J. Immune checkpoint inhibitors in advanced cutaneous squamous cell carcinoma: A systemic review and meta-analysis. Skin Res Technol 29, e13229 (2023).

34. Que, S.K.T., Zwald, F.O. & Schmults, C.D. Cutaneous squamous cell carcinoma: Incidence, risk factors, diagnosis, and staging. Journal of the American Academy of Dermatology 78, 237–247 (2018).

35. Chang, A.L.S., Zaba, L. & Kwong, B.Y. Immunotherapy for keratinocyte cancers. Part II: Identification and management of cutaneous side effects of immunotherapy treatments. Journal of the American Academy of Dermatology 88, 1243–1255 (2023).

36. Leoni, G. et al. A Genetic Vaccine Encoding Shared Cancer Neoantigens to Treat Tumors with Microsatellite Instability. Cancer Research 80, 3972–3982 (2020).

37. Li, F. et al. Neoantigen vaccination induces clinical and immunologic responses in non-small cell lung cancer patients harboring EGFR mutations. J Immunother Cancer 9 (2021).

38. Adams, A.C. et al. Solar Simulated Light Induces Cutaneous Squamous Cell Carcinoma in Inbred Mice: A Clinically Relevant Model to Investigate T-Cell Responses. J Invest Dermatol 141, 2990–2993.e2996 (2021).

39. Freeman, A. et al. Comparative immune phenotypic analysis of cutaneous Squamous Cell Carcinoma and Intraepidermal Carcinoma in immune-competent individuals: proportional representation of CD8+ T-cells but not FoxP3+ Regulatory T-cells is associated with disease stage. PLoS One 9, e110928 (2014).

40. Zeng, Z. et al. IFN-γ Critically Enables the Intratumoural Infiltration of CXCR3(+) CD8(+) T Cells to Drive Squamous Cell Carcinoma Regression. Cancers (Basel) 13 (2021).

41. Nasti, T.H. et al. Differential roles of T-cell subsets in regulation of ultraviolet radiation induced cutaneous photocarcinogenesis. Photochem Photobiol 87, 387–398 (2011).

42. Yusuf, N. et al. Antagonistic roles of CD4+ and CD8+ T-cells in 7,12-dimethylbenz(a)anthracene cutaneous carcinogenesis. Cancer Res 68, 3924–3930 (2008).

43. Dodagatta-Marri, E. et al. α-PD-1 therapy elevates Treg/Th balance and increases tumor cell pSmad3 that are both targeted by α-TGFβ antibody to promote durable rejection and immunity in squamous cell carcinomas. J Immunother Cancer 7, 62 (2019).

44. Cohen, C.J. et al. Isolation of neoantigen-specific T cells from tumor and peripheral lymphocytes. J Clin Invest 125, 3981–3991 (2015).

45. Strønen, E. et al. Targeting of cancer neoantigens with donor-derived T cell receptor repertoires. Science 352, 1337–1341 (2016).

46. Capietto, A.H. et al. Mutation position is an important determinant for predicting cancer neoantigens. J Exp Med 217 (2020).

47. Ebrahimi-Nik, H. et al. Reversion analysis reveals the in vivo immunogenicity of a poorly MHC I-binding cancer neoepitope. Nat Commun 12, 6423 (2021).

48. Devlin, J.R. et al. Structural dissimilarity from self drives neoepitope escape from immune tolerance. Nat Chem Biol 16, 1269–1276 (2020).

49. Borbulevych, O.Y. et al. Structures of MART-126/27-35 Peptide/HLA-A2 complexes reveal a remarkable disconnect between antigen structural homology and T cell recognition. J Mol Biol 372, 1123–1136 (2007).

50. Anderson, K.S., Alexander, J., Wei, M. & Cresswell, P. Intracellular transport of class I MHC molecules in antigen processing mutant cell lines. J Immunol 151, 3407–3419 (1993).

51. Asmann, Y.W. et al. TREAT: a bioinformatics tool for variant annotations and visualizations in targeted and exome sequencing data. Bioinformatics 28, 277–278 (2012).

52. de Bruijn, I. et al. Analysis and Visualization of Longitudinal Genomic and Clinical Data from the AACR Project GENIE Biopharma Collaborative in cBioPortal. Cancer Res 83, 3861–3867 (2023).

53. Shukla, S.A. et al. Comprehensive analysis of cancer-associated somatic mutations in class I HLA genes. Nat Biotechnol 33, 1152–1158 (2015).

54. Maiers, M., Gragert, L. & Klitz, W. High-resolution HLA alleles and haplotypes in the United States population. Hum Immunol 68, 779–788 (2007).

55. Ye, J. et al. Primer-BLAST: a tool to design target-specific primers for polymerase chain reaction. BMC Bioinformatics 13, 134 (2012).

56. Ji, A.L. et al. Multimodal Analysis of Composition and Spatial Architecture in Human Squamous Cell Carcinoma. Cell 182, 497–514.e422 (2020).

57. Chitsazzadeh, V. et al. Cross-species identification of genomic drivers of squamous cell carcinoma development across preneoplastic intermediates. Nat Commun 7, 12601 (2016).

58. Auton, A. et al. A global reference for human genetic variation. Nature 526, 68–74 (2015).

59. Li, H. Aligning sequence reads, clone sequences and assembly contigs with BWA-MEM. arXiv: Genomics (2013).

60. Li, H. et al. The Sequence Alignment/Map format and SAMtools. Bioinformatics 25, 2078–2079 (2009).

61. Van der Auwera, G.A. & O’Connor, B.D. Genomics in the cloud : using Docker, GATK, and WDL in Terra, First edition. edn. O’Reilly Media: Sebastopol, CA, 2020.

62. Saunders, C.T. et al. Strelka: accurate somatic small-variant calling from sequenced tumor-normal sample pairs. Bioinformatics 28, 1811–1817 (2012).

63. McLaren, W. et al. The Ensembl Variant Effect Predictor. Genome Biol 17, 122 (2016).

64. Hundal, J. et al. pVAC-Seq: A genome-guided in silico approach to identifying tumor neoantigens. Genome Med 8, 11 (2016).

65. Fairley, S., Lowy-Gallego, E., Perry, E. & Flicek, P. The International Genome Sample Resource (IGSR) collection of open human genomic variation resources. Nucleic Acids Res 48, D941–d947 (2020).

66. Cho, Y., Gorina, S., Jeffrey, P.D. & Pavletich, N.P. Crystal structure of a p53 tumor suppressor-DNA complex: understanding tumorigenic mutations. Science 265, 346–355 (1994).

67. Sehnal, D. et al. Mol* Viewer: modern web app for 3D visualization and analysis of large biomolecular structures. Nucleic Acids Research 49, W431–W437 (2021).

68. Jumper, J. et al. Highly accurate protein structure prediction with AlphaFold. Nature 596, 583–589 (2021).

69. Scott, D.M., Ehrmann, I.E., Ellis, P.S., Chandler, P.R. & Simpson, E. Why do some females reject males? The molecular basis for male-specific graft rejection. J Mol Med (Berl) 75, 103–114 (1997).

70. Tomayko, M.M. & Reynolds, C.P. Determination of subcutaneous tumor size in athymic (nude) mice. Cancer Chemother Pharmacol 24, 148–154 (1989).

71. Patro, R., Duggal, G., Love, M.I., Irizarry, R.A. & Kingsford, C. Salmon provides fast and bias-aware quantification of transcript expression. Nat Methods 14, 417–419 (2017).

72. Ye, J. et al. Primer-BLAST: A tool to design target-specific primers for polymerase chain reaction. BMC Bioinformatics 13, 134 (2012).

73. Phipps-Yonas, H., Semik, V. & Hastings, K.T. GILT expression in B cells diminishes cathepsin S steady-state protein expression and activity. Eur J Immunol 43, 65–74 (2013).

